# Transcriptomic and spatial GABAergic neuron subtypes in zona incerta mediate distinct innate behaviors

**DOI:** 10.1101/2025.02.10.637400

**Authors:** Mengyue Zhu, Jieqiao Peng, Mi Wang, Shan Lin, Huiying Zhang, Yu Zhou, Xinyue Dai, Huiying Zhao, Yan-qin Yu, Li Shen, Xiao-Ming Li, Jiadong Chen

## Abstract

Understanding the anatomical connection and behaviors of transcriptomic neuron subtypes is critical to delineating cell type-specific functions in the brain. Here we integrated single-nucleus transcriptomic sequencing, *in vivo* circuit mapping, optogenetic and chemogenetic approaches to dissect the molecular identity and function of heterogeneous GABAergic neuron populations in the zona incerta (ZI) in mice, a region involved in modulating various behaviors. By microdissecting ZI for transcriptomic and spatial gene expression analyses, our results revealed two non-overlapping *Ecel1-* and *Pde11a*-expressing GABAergic neurons with dominant expression in the rostral and medial zona incerta (ZIr^Ecel1^ and ZIm^Pde11a^), respectively. The GABAergic projection from ZIr^Ecel1^ to periaqueductal gray mediates self-grooming, while the GABAergic projection from ZIm^Pde11a^ to the oral part of pontine reticular formation promotes transition from sleep to wakefulness. Together, our results revealed the molecular markers, spatial organization and specific neuronal circuits of two discrete GABAergic projection neuron populations in segregated subregions of the ZI that mediate distinct innate behaviors, advancing our understanding of the functional organization of the brain.

## Introduction

The molecular, morphological and anatomical features of neurons imply distinct neuronal subtypes, axonal projection pattern, and behavioral functions in the central nervous system^1, 2^. Recent advances in single-cell transcriptomics reveal cell type specific genes expression and molecular markers of transcriptomic cell subtypes^3–5^. Extensive efforts have been applied to classify neuron subtypes by morphological and electrophysiological characteristics^6–8^, in conjunction with single-cell transcriptomic analysis^9–13^. Spatial transcriptomic analysis discovered diverse cell subtypes and their anatomical organization across the whole brain^14–16^. In addition, the integrated retrograde or anterograde transsynaptic tracing with spatial transcriptomic analyses revealed the long-range synaptic connectivity of neuronal subtypes, connecting transcriptomic clusters with axonal projections at scale^4, 17–20^. However, the behavioral functions of different transcriptomic neuronal subtypes are yet to be determined in behaving animals. Therefore, investigating the anatomical connection and behavior of transcriptomic and spatial neuron subtypes is important to reveal the functional organization of the brain.

The GABAergic projection neurons establish long-range efferent projections and may promote efficient information transfer between distant brain networks compared with the well-studied cortical interneurons ^21, 22^. Recent studies revealed the presence of long-range GABAergic projection neurons in various brain regions including the prefrontal cortex^23, 24^, entorhinal cortex^25^, hippocampus^25, 26^, and also top-down afferents from the zona incerta (ZI)^27^, basal ganglia^28^, and septum^29^, but their sparse distribution makes it difficult to identify the molecular markers for this heterogeneous population by conventional single-cell RNA sequencing approach^21, 22^. The zona incerta (ZI), located in the ventral thalamus, predominantly comprises GABAergic neurons that project to distant brain regions and mediate various behavioral functions^30, 31^. The heterogeneous neuron populations in the ZI can be divided into cytoarchitectonic subregions or sectors along the anteroposterior axis, which form extensive connections with multiple brain regions and may function as one of the integrative nodes in the brain^30–33^. Recent studies in rodents and non-human primates revealed various behavioral functions upon stimulating subregions of the ZI ^34–42^. However, the lack of molecular markers makes it difficult to selectively target or to investigate the function of GABAergic projection neurons in different subregions of the ZI^21, 22^.

In the present study, by microdissecting the ZI for single-nucleus transcriptomic sequencing, we investigated the molecular subtypes and spatial localization of GABAergic neuron populations in different subregions of the ZI. Through constructing transgenic mice to label region-specific GABAergic neuron subtypes in ZI, integrated with *in vivo* circuit mapping, and optogenetic and chemogenetic manipulation, we revealed two discrete subpopulations of GABAergic projection neurons in segregated subregions of the ZI that exert distinct behavioral functions.

## Results

### Single-nuclei transcriptomic profiling revealed distinct cell clusters and region specific gene expression in the zona incerta

To analyze the molecular diversity of cell types in ZI, we micro-dissected the brain region according to the distribution of fluorescent reporter from postnatal day 40 (P40) Gad67-GFP transgenic mice in ZI and the anatomy from the reference mouse brain atlas of Allen Mouse Brain Common Coordinate Framework^33^. We dissociated the micro-dissected brain tissue into single nuclei suspension and analyzed the single-nuclei transcriptome of ZI in the adult mouse brain by 10x genomics. We harvested 15,375 single nuclei from two independent cohorts of 10x genomics single-nucleus RNA sequencing experiments and obtained similar cell clusters by reciprocal principle component analysis (RPCA) (Fig. 1a and Supplementary Fig. 1a-c). The cell clusters were displayed by uniform manifold approximation and projection (UMAP) and annotated with canonical cell type specific marker genes. We identified cell clusters including major cell types such as inhibitory neurons (*Rbfox3*, *Gad2*), excitatory neurons (*Rbfox3*, *Slc17a6*)^43–45^, dopaminergic neurons (*Slc6a3*)^46, 47^, microglia (*Cx3cr1*)^44, 48^, astrocytes (*Aldh1a1*)^49^, pericytes (*Pdgfrb*)^44^, endothelial cells (*Cldn5*)^46, 47^, oligodendrocyte precursor cells (OPCs, *Pdgfra*), differentiation-committed oligodendrocyte precursors (COPs, *Tns3*), oligodendrocytes (*Mog*) and mature oligodendrocytes (MOLs, *Apod*) ^50^(Fig. 1a and Supplementary Fig. 1d).

**Fig. 1:**
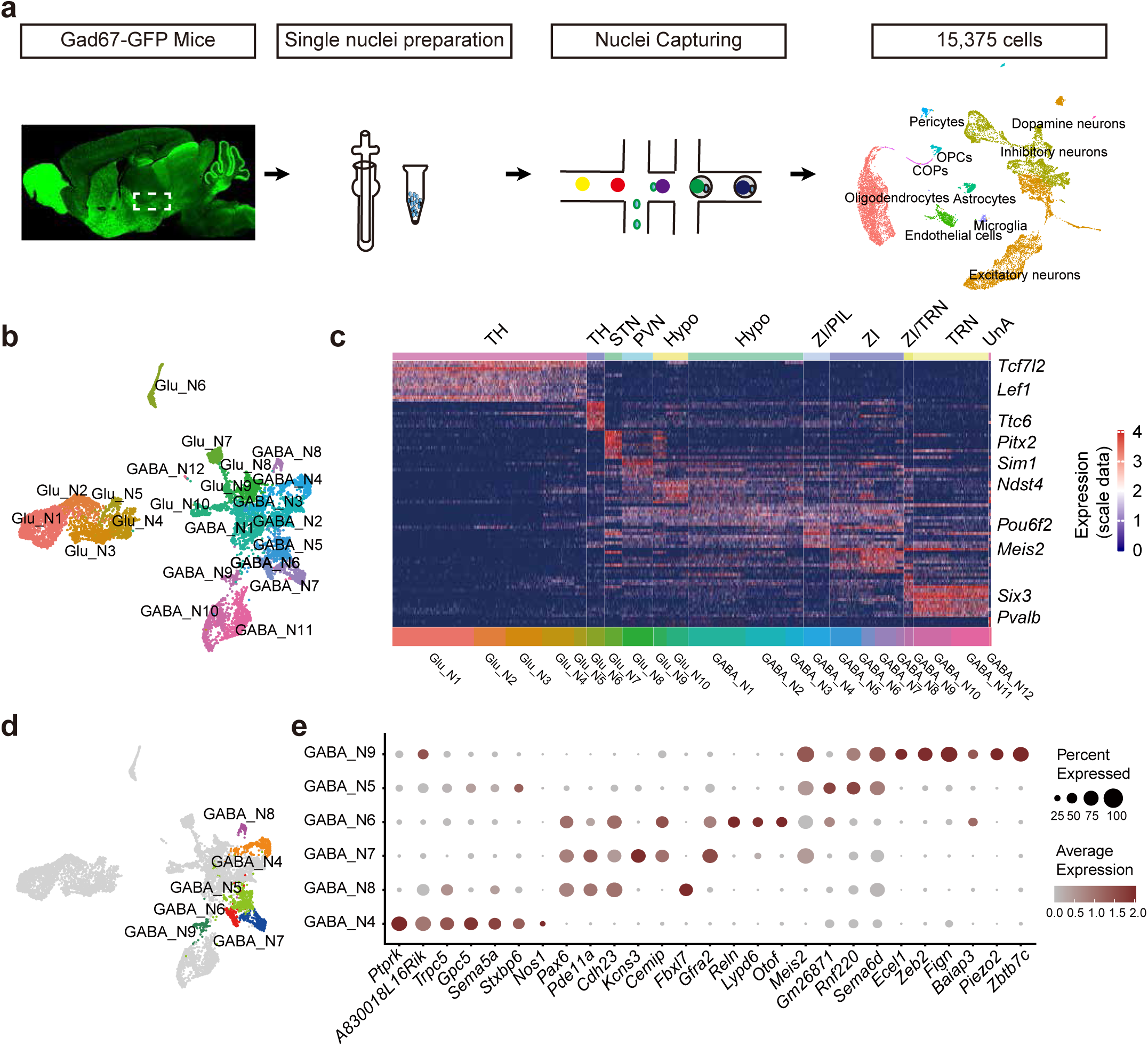
Single-nucleus transcriptomic sequencing reveals major cell type composition in zona incerta. **a,** Schematic diagram showing the workflow for microdissection of zona incerta from brain slices in green fluorescent protein knock-in transporter line Gad67-GFP mice, single nucleus isolation and RNA sequencing by 10x Genomics. UMAP plot of the processed dataset containing 15,375 single nuclei collected from two independent cohorts of 10x Genomics experiments (n = 6 mice for each cohort). The cell clusters were annotated according to canonical cell type-specific marker genes. **b,** UMAP plot showing re-clustering the excitatory and inhibitory neurons into ten glutamatergic neuron clusters (Glu_N1 to Glu_N10) and twelve GABAergic neuron clusters (GABA_N1 to GABA_N12). **c,** Heatmap showing the differentially expressed genes (DEGs) among neuron clusters from snRNA-seq dataset. The regional identity (top) of the neuron clusters were annotated by the expression of region specific DEGs (right). TH, Thalamus; STN, Subthalamus nucleus; PVN, Paraventricular nucleus; Hypo, Hypothalamus; ZI, Zona incerta; PIL, Posterior intralaminar thalamus nucleus; TRN, Thalamic reticular nucleus; UnA, Un-Annotated region. **d,** UMAP plot showing the GABAergic neuron clusters (GABA_N4 to GABA_N9) associated with zona incerta. **e,** Dotplot showed the differential expressed genes (DEGs) of GABAergic neuron clusters associated with zona incerta. Source data are provided as a Source Data file.

We further reclustered the neuronal populations and revealed 10 clusters of glutamatergic neurons (Glu_N1 – Glu_N10) and 12 clusters of GABAergic neurons (GABA_N1 – GABA_N12) that showed distinct differentially expressed genes (DEGs) among neuron subtypes in ZI and neighboring brain regions (Fig. 1b, c). Based on *in situ* hybridization of gene expression in the adult mouse brain from Allen brain atlas (https://mouse.brain-map.org/)^51^, we annotated the anatomical locations of neuronal clusters and identified marker genes for ZI and neighboring brain regions in the subcortical forebrain: ZI (*Pax6, Meis2*), thalamus (*Tcf7l2, Lef1*) located dorsal to ZI, subthalamus nucleus (*Ttc6, Pitx2*) located ventral to ZI, paraventricular nucleus (*Sim1*, *Ndst4*) located ventral-medial to ZI, and reticular nucleus (*Six3*, *Pvalb*) located dorsal-lateral to ZI (Supplementary Fig. 2a).

GABAergic neurons in ZI projected to multiple brain regions and were implicated in various behaviors, however, the molecular profiles of different GABAergic neuron subtypes remain incompletely understood. Our data revealed six GABAergic neuron clusters in ZI (GABA_N4 to GABA_N9, Fig. 1d, e; Supplementary Fig. 2b, c and Supplementary table 1). Differential gene expression analysis revealed marker genes for different GABAergic neuron clusters that exhibited different anatomical distribution pattern in ZI, as demonstrated by *in situ* hybridization of gene expression in the Allen Mouse Brain Atlas (https://mouse.brain-map.org/)^51^. The GABA_N9 neuron cluster highly expressed *Ecel1* (endothelin converting enzyme like 1). The GABA_N6, GABA_N7 and GABA_N8 neuron clusters shared differentially expressed genes and highly expressed *Cdh23* (cadherin related 23), *Pax6* (paired box 6) and *Pde11a* (phosphodiesterase 11A). The GABA_N4 neuron cluster highly expressed *Ptprk* (protein tyrosine phosphatase receptor type K) and *Nos1* (nitric oxide synthase 1) (Fig. 2a and Supplementary Fig. 2c). These results revealed distinct transcriptomic profiles and candidate marker genes for GABAergic neurons in the ZI by single-nucleus RNA sequencing analysis.

**Fig. 2:**
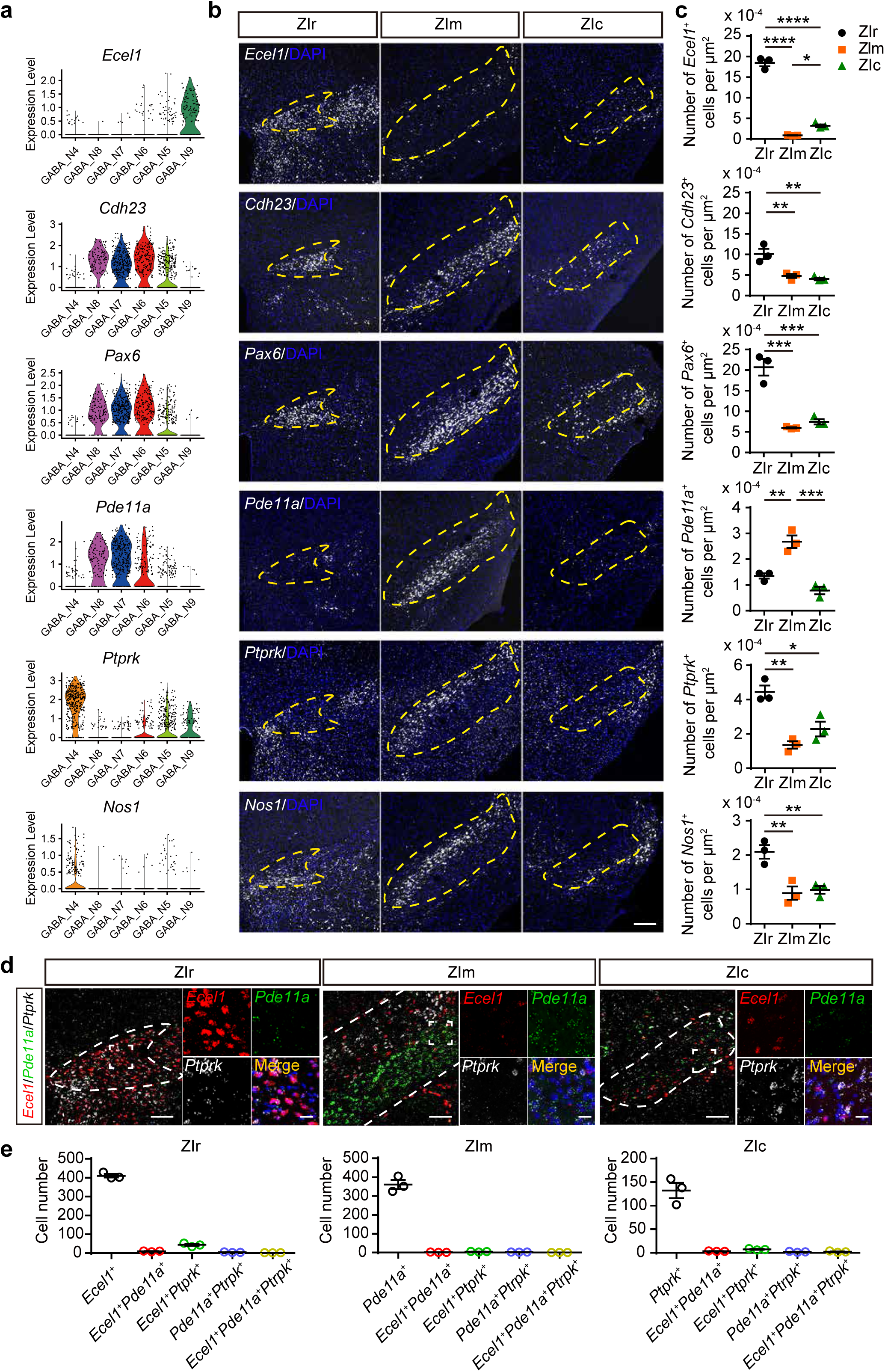
Region-specific genes expression of GABAergic neuron subtypes in subregions of zona incerta. **a,** Violin Plots showed the expression level of selected DEG genes of GABAergic neuron clusters in zona incerta. **b,** Representative images (**b**) and statistical results (**c**) showing the density of *Ecel1* (ZIr *vs* ZIm, \*\*\*\**P* < 0.0001; ZIr *vs* ZIc, \*\*\*\**P* < 0.0001; ZIm *vs* ZIc, \**P* = 0.0443), *Cdh23* (ZIr *vs* ZIm, \*\**P* =0.0075; ZIr *vs* ZIc, \*\**P* =0.0039)*, Pax6* (ZIr *vs* ZIm, \*\*\**P* =0.0004; ZIr *vs* ZIc, \*\*\**P* =0.0007), *Pde11a* (ZIr *vs* ZIm, \*\**P* =0.0038; ZIm *vs* ZIc, \*\*\**P* =0.0006), *Ptprk* (ZIr *vs* ZIm, \*\**P* =0.0020; ZIr *vs* ZIc, \**P* =0.0119), *Nos1* (ZIr *vs* ZIm, \*\**P* =0.0062; ZIr *vs* ZIc, \*\**P* =0.0092) in the ZIr, ZIm and ZIc, n = 9 brain slices from 3 mice per group. Scale bar, 200 μm. One-way ANOVA and tukey’s multiple comparisons test. **d,** Representative images (**d**) and statistical results (**e**) showing the overlap between *Ecel1*, *Pde11a* and *Ptprk* in different brain sections of ZIr, ZIm and ZIc. The cell numbers in ZIr (*Ecel1^+^*: 410.0 ± 9.539 cells; *Ecel1^+^Pde11a*^+^: 9.33 ± 1.20 cells; *Ecel1^+^Ptprk*^+^: 43.33 ± 5.49 cells; *Pde11a^+^Ptprk*^+^: 4.33 ± 0.88 cells; *Ecel1^+^Pde11a^+^Ptprk*^+^: 1.67 ± 0.33 cells), ZIm (*Pde11a*^+^: 361.0 ± 23.81 cells; *Ecel1^+^Pde11a*^+^: 1.67 ± 0.67 cells; *Ecel1^+^Ptprk*^+^: 4.67 ± 0.88 cells; *Pde11a^+^Ptprk*^+^: 2.67 ± 0.33 cells; *Ecel1^+^Pde11a^+^Ptprk*^+^: 0 ± 0 cells) and ZIc (*Ptprk*^+^: 132.3 ± 16.13 cells; *Ecel1^+^Pde11a*^+^: 3.33 ± 0.33 cells; *Ecel1^+^Ptprk*^+^: 7.33 ± 0.88 cells; *Pde11a^+^Ptprk*^+^: 2.00 ± 0.58 cells; *Ecel1^+^Pde11a^+^Ptprk*^+^: 2.33 ± 0.88 cells) were shown respectively. N = 9 brain slices from 3 mice per group. Scale bars, 100 μm and 20 μm (zoom-in image). Data are presented as mean ± SEM. Source data are provided as a Source Data file.

### Candidate genes for GABAergic neuron subtypes in subregions of zona incerta

We validated the expression and spatial distribution of the above six GABAergic marker genes in different anatomical subregions of ZI along the anteroposterior axis by fluorescence *in situ* hybridization (FISH). We found that *Ecel1* exhibited highest distribution in the rostral ZI (ZIr: 59.56 ± 0.93%; ZIm: 14.41 ± 1.33%; ZIc: 26.03 ± 1.76%) compared with other marker genes expression in the ZIr (*Cdh23*: 22.37 ± 1.40%; *Pax6*: 29.20 ± 1.83%; *Nos1*: 22.98 ± 1.68%; *Ptprk*: 25.59 ± 1.67%). The marker gene *Pde11a* exhibited highest and most selective distribution in the medial ZI (ZIm: 80.49 ± 1.57%; ZIr: 7.873 ± 0.55%; ZIc: 11.63 ± 2.04%) compared with other marker genes expression in the ZIm (*Cdh23*: 54.60 ± 0.89%; *Pax6*: 43.97 ± 1.79%; *Nos1*: 49.20 ± 5.38%). We found that 33.82 ± 6.63% of the total *Ptprk* positive cells in the ZI were distributed in the caudal ZI, but showed no significant difference between different subregions of ZI (ZIr: 25.59 ± 1.67%; ZIm: 40.59 ± 6.35%. Fig. 2b and Supplementary Fig. 2d).

We further demonstrated that the density of *Ecel1^+^* cells is relatively higher than *Pde11a*^+^ or *Ptprk*^+^ cells in the ZIr (*Ecel1^+^*: 1848 ± 82.52 per mm^2^; *Pde11a*^+^: 134.8 ± 10.60 per mm^2^; *Ptprk*^+^: 444.9 ± 38.15 per mm^2^), and showed few overlap with *Pde11a* and *Ptprk* (*Ecel1^+^*: 410.0 ± 9.539 cells; *Ecel1^+^Pde11a*^+^: 9.33 ± 1.20 cells; *Ecel1^+^Ptprk*^+^: 43.33 ± 5.49 cells; *Pde11a^+^Ptprk*^+^: 4.33 ± 0.88 cells; *Ecel1^+^Pde11a^+^Ptprk*^+^: 1.67 ± 0.33 cells). The *Pde11a*^+^ cells show higher density in the ZIm (*Ecel1^+^*: 85.77 ± 4.066 per mm^2^; *Pde11a*^+^: 268.2 ± 24.12 per mm^2^; *Ptprk*^+^: 136.2 ± 21.35 per mm^2^) and also show very few overlap with *Ecel1* and *Ptprk* (*Pde11a*^+^: 361.0 ± 23.81 cells; *Ecel1^+^Pde11a*^+^: 1.67 ± 0.67 cells; *Ecel1^+^Ptprk*^+^: 4.67 ± 0.88 cells; *Pde11a^+^Ptprk*^+^: 2.67 ± 0.33 cells; *Ecel1^+^Pde11a^+^Ptprk*^+^: 0 ± 0 cells). In contrast, the density of *Ptprk*^+^ cells showed little differences with *Ecel1^+^* or *Pde11a*^+^ cells in the ZIc (*Ecel1^+^*: 317.8 ± 34.96 per mm^2^; *Pde11a*^+^: 78.27 ± 14.38 per mm^2^; *Ptprk*^+^: 228.9 ± 42.81 per mm^2^. Fig. 2c - e) and *Ptprk*^+^ cells rarely overlap with *Ecel1^+^* or *Pde11a*^+^ cells (*Ptprk*^+^: 132.3 ± 16.13 cells; *Ecel1^+^Pde11a*^+^: 3.33 ± 0.33 cells; *Ecel1^+^Ptprk*^+^: 7.33 ± 0.88 cells; *Pde11a^+^Ptprk*^+^: 2.00 ± 0.58 cells; *Ecel1^+^Pde11a^+^Ptprk*^+^: 2.33 ± 0.88 cells). These results demonstrated that the expression of *Ecel1* and *Pde11a* are non-overlapping across different subregions of the ZI along the anteroposterior axis (Fig. 2d, e). Together, these results revealed that *Ecel1-* and *Pde11a-*expressing GABAergic neurons are non-overlapping and exhibit dominant expression in the rostral and medial ZI, respectively.

### Ecel1-positive neurons in the rostral ZI and Pde11a-positive neurons in the medial ZI displayed distinct efferent projections

Previous studies showed that GABAergic neurons in the ZI projected to different brain regions and exhibited various behavioral functions^30, 31^. We hypothesized that GABAergic projection neurons in different subregions of the ZI may drive distinct functions. Firstly, we further confirmed that *Ecel1* and *Pde11a* are co-expressed with GABAergic neuronal marker *Slc32a1* (Vgat, vesicular GABA transporter) in the ZIr (*Ecel1^+^Slc32a1^+^ /Ecel1^+^:* 96.39 ± 1.19%; *Ecel1^+^Slc32a1^+^/Slc32a1^+^*: 59.69 ± 6.28%) and the ZIm (*Pde11a^+^Slc32a1^+^/Pde11a^+^*: 100 ± 0.00%*; Pde11a^+^Slc32a1^+^/Slc32a1^+^*: 33.92 ± 9.67%, n = 9 brain slices from 3 mice per group. Fig. 3a-c). And for selectively target of GABAergic projection neuron subtypes in the rostral or medial ZI, we constructed two transgenic mice to selectively label *Ecel1*-expressing neurons (Ecel1-Cre) and *Pde11a*-expressing neurons (Pde11a-Cre), respectively (Fig. 3d). We verified the specificity of transgenic mice by examining the co-expression of Cre recombinase with *Ecel1* or *Pde11a*. The Cre recombinase expression in Ecel1-Cre is predominantly distributed in the ZIr (71.95 ± 1.12%, n = 9 brain slices from 3 mice) and highly co-expressed with *Ecel1* in ZIr (93.87 ± 1.45% *Ecel1*^+^Cre^+^ double positive neurons out of Cre^+^ neurons. 96.41 ± 0.92% *Ecel1*^+^Cre^+^ double positive neurons out of *Ecel1*^+^ neurons). In contrast, the expression of Cre recombinase in Pde11a-Cre mice is predominantly distributed in the ZIm (90.33 ± 2.91%, n = 9 brain slices from 3 mice) and highly co-expressed with *Pde11a* (99.70 ± 0.15% *Pde11a*^+^Cre^+^ double positive neurons out of Cre^+^ neurons, 98.91 ± 0.22% *Pde11a*^+^Cre^+^ double positive neurons out of *Pde11a*^+^ neurons. Fig. 3e-i). These results demonstrated high fidelity of selective labeling of *Ecel1* and *Pde11a*-expressing neurons by the Ecel1-Cre and Pde11a-Cre transgenic mice.

**Fig. 3:**
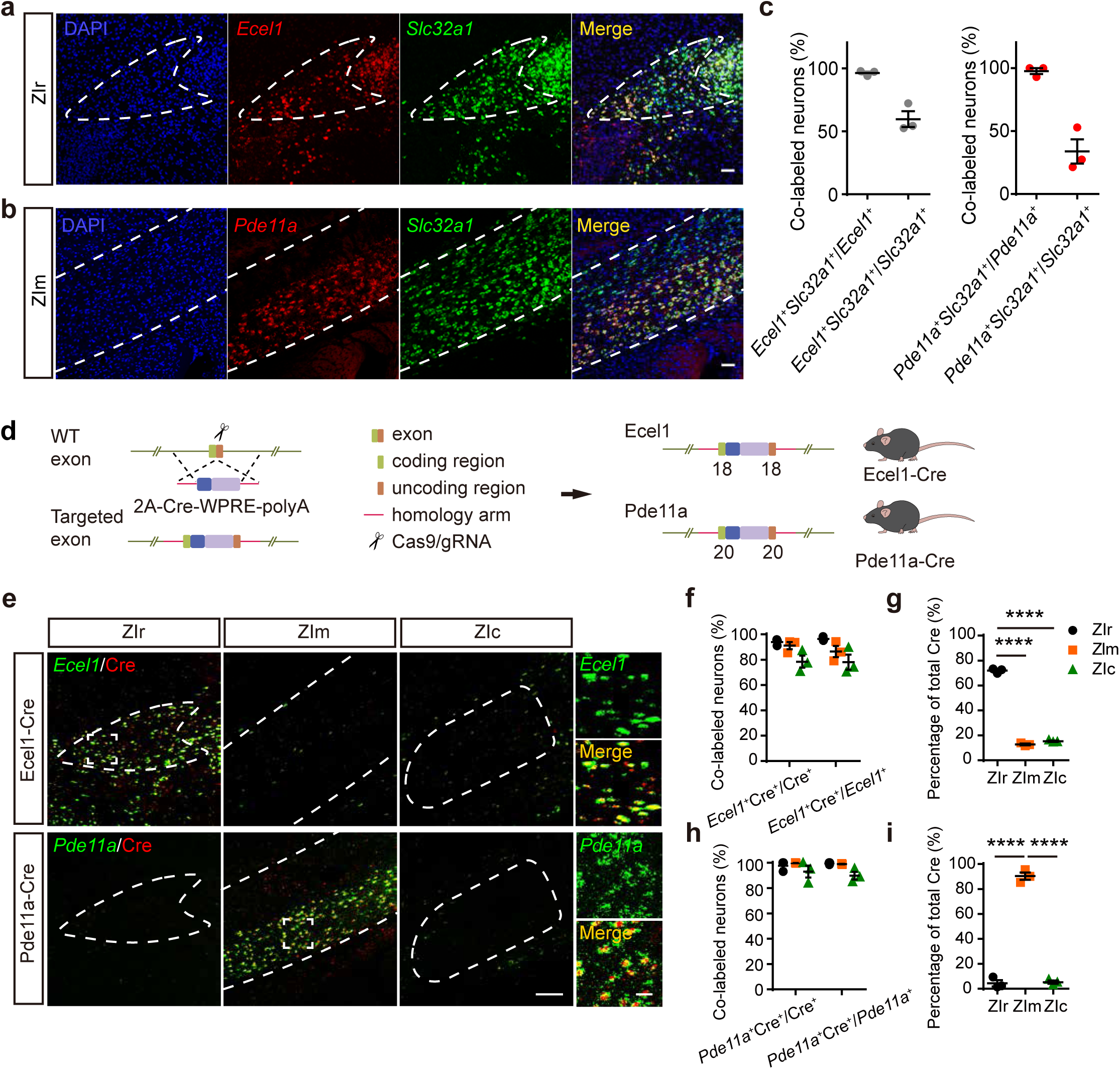
Generation and validation of transgenic mice Ecel1-Cre and Pde11a-Cre. **a,** Representative images showing the expression of *Ecel1* (**a**) and *Pde11a* (**b**) in *Slc32a1* (Vgat) positive neurons in ZIr or ZIm. Scale bar, 50μm. **c,** The percentage of *Ecel1^+^Slc32a1^+^* cells relative to all *Ecel1^+^* or *Slc32a1^+^* cells in ZIr (*Ecel1^+^Slc32a1^+^/Ecel1^+^*, 96.39±1.19%; *Ecel1^+^Slc32a1^+^/Slc32a1^+^*, 59.69±6.28%) and the percentage of *Pde11a^+^Slc32a1^+^* cells relative to all *Pde11a^+^*or *Slc32a1^+^* cells in ZIm (*Pde11a^+^Slc32a1^+^/Pde11a^+^*, 97.7±2.3%; *Pde11a^+^Slc32a1^+^/Slc32a1^+^*, 33.92±9.67%). n=9 brain slices from 3 mice per group. **d,** Schematic diagram showing the generation of Ecel1-Cre or Pde11a-Cre knock-in mice. **e,** Representative images showing the Cre and *Ecel1* or the Cre and *Pde11a* expression in ZIr, ZIm and ZIc. Scale bars, 100μm and 20μm (zoom-in image). **f,** The percentage of *Ecel1^+^*Cre^+^ relative to all Cre^+^ or *Ecel1*^+^ neurons in ZIr (*Ecel1^+^*Cre^+^/Cre^+^,93.87±1.45%; *Ecel1^+^*Cre^+^/*Ecel1*^+^, 96.41±0.92%), Zim (*Ecel1^+^*Cre^+^/Cre^+^, 91.11±2.81%; *Ecel1^+^*Cre^+^/*Ecel1*^+^, 86.46±4.39%) and Zic (*Ecel1^+^*Cre^+^/Cre^+^, 78.41±4.75%; *Ecel1^+^*Cre^+^/*Ecel1*^+^, 78.00±5.95%) of Ecel1-Cre mice. n = 9 brain slices from 3 mice per group. **g,** Statistical results showing the expression and distribution of Cre in the ZIr (71.95±1.12%), ZIm (12.76±0.67%) and ZIc (15.28±0.49%) of Ecel1-Cre mice. ZIr *vs* ZIm, \*\*\*\**P*<0.0001; ZIr *vs* ZIc, \*\*\*\**P*<0.0001. N = 9 brain slices from 3 mice per group. **h,** The percentage of *Pde11a^+^*Cre^+^ relative to all Cre+ or *Pde11a^+^* neurons in ZIr (*Pde11a^+^*Cre^+^/Cre^+^, 97.70±2.30%; *Pde11a^+^*Cre^+^/*Pde11a^+^*, 99.48±0.52%), Zim (*Pde11a^+^*Cre^+^/Cre^+^, 99.70±0.15%; *Pde11a^+^*Cre^+^/*Pde11a^+^*, 98.91±0.22%) and Zic (*Pde11a^+^*Cre^+^/Cre^+^, 93.00±4.72%; *Pde11a^+^*Cre^+^/*Pde11a^+^*, 89.73±3.06%) of Pde11a-Cre mice. n=9 brain slices from 3 mice per group. **i,** Statistical results showing the expression and distribution of Cre in the ZIr (4.43±2.54%), ZIm (90.33±2.91%) and ZIc (5.24±1.27%) of Pde11a-Cre mice. ZIr *vs* ZIm, \*\*\*\**P*<0.0001; ZIm *vs* ZIc, \*\*\*\**P*< 0.0001. N=9 brain slices from 3 mice per group. One-way ANOVA and tukey’s multiple comparisons test for (**g**) and (**i**). Data are presented as mean±SEM. Source data are provided as a Source Data file.

Next, we examined the axonal projections of ZIr^Ecel1^ and ZIm^Pde11a^ GABAergic neurons by stereotaxic injection of AAV virus carrying Cre-dependent expression of membrane-bound GFP (mGFP) and synaptophysin-mRuby (SYP-mRuby) (Fig. 4a, b and Supplementary Fig. 3a, b). We found the axon terminals of ZIr^Ecel1^ GABAergic projection neurons distributed in several brain regions including periaqueductal gray (PAG), lateral septal nucleus (LS), preoptic area (POA), paraventricular nucleus of the thalamus (PVT), reuniens thalamic nucleus (Re), lateral hypothalamic area (LH), mediodorsal thalamic nucleus (MD) (Fig. 4c-e and Supplementary Fig. 3c). In contrast, the axon terminals of ZIm^Pde11a^ GABAergic neurons distributed in brain regions including pontine reticular formation (PRF), posterior complex of the thalamus (PO), auditory cortex (AuV), anterior pretectal nucleus, dorsal part (APTD), nucleus of Darkschewitsch (Dk), red nucleus (RN) and superior colliculus (SC) (Fig. 4c-e and Supplementary Fig. 3c). These results demonstrated divergent efferent projections of ZIr^Ecel1^ and ZIm^Pde11a^ GABAergic neurons to distinct downstream brain regions. For example, the periaqueductal gray (PAG) received extensive axonal projections from ZIr^Ecel1^ neurons (ZIr^Ecel1^-PAG pathway), but very few projections from ZIm^Pde11a^ neurons. In contrast, pontine reticular formation (PRF) received axonal projections predominantly from ZIm^Pde11a^ (ZIm^Pde11a^-PRF pathway), but not ZIr^Ecel1^ GABAergic neurons (Fig. 4c, d). Therefore, the efferent projection of ZIr^Ecel1^ and ZIm^Pde11a^ GABAergic neurons to different downstream target brain regions may imply their distinct behavioral functions.

**Fig. 4:**
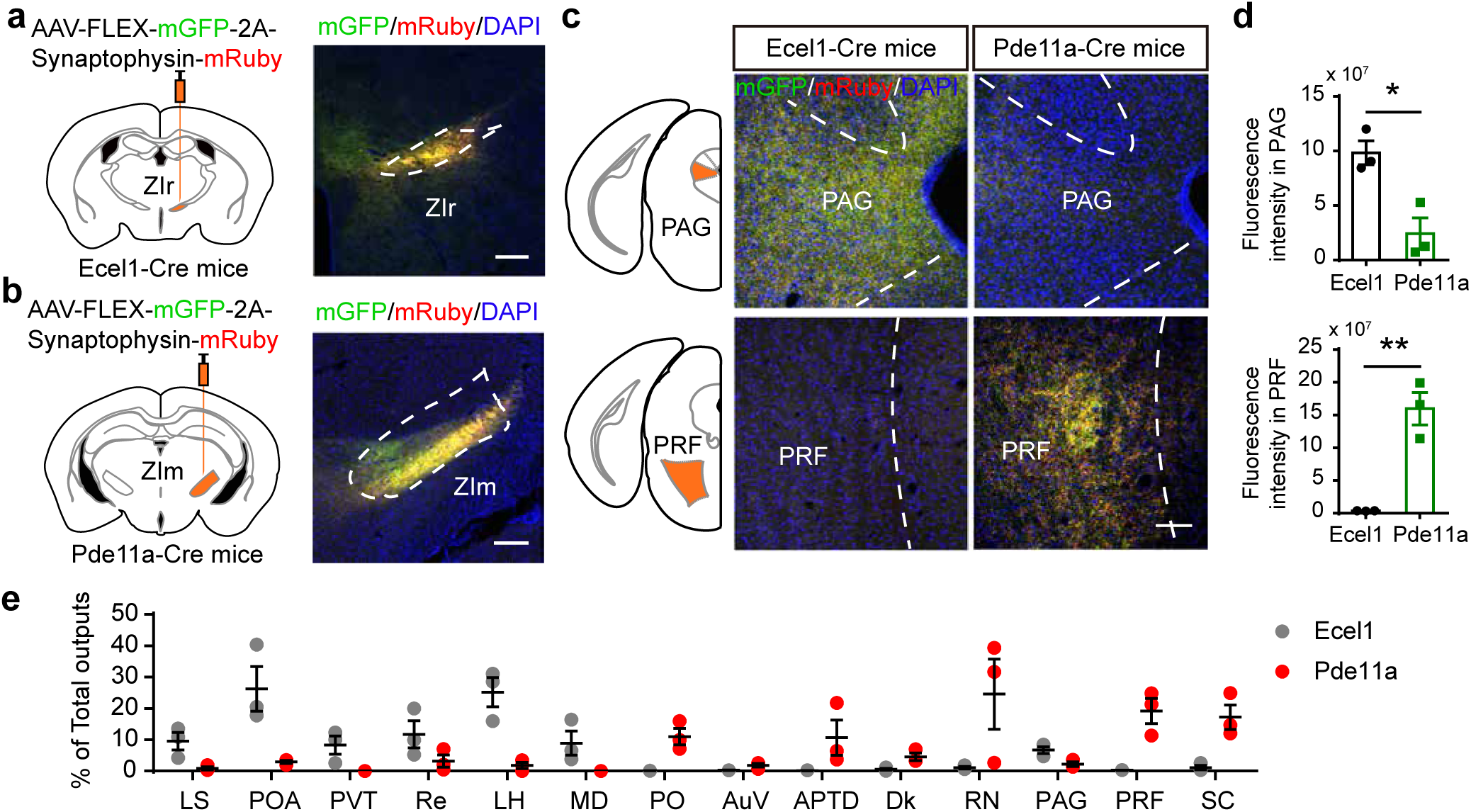
Distinct efferent projections of Ecel1 positive neurons in the rostral ZI and Pde11a positive neurons in the medial ZI. **a,** Schematic diagram and representative images showing the stereotaxic unilateral injection of Cre-dependent AAV mediated mGFP and synaptophysin-mRuby to specifically label Ecel1 positive neurons in the rostral ZI (ZIr^Ecel1^) of Ecel1-Cre mice (**a**) or Pde11a positive neurons in the medial ZI (ZIm^Pde11a^) of Pde11a-Cre mice (**b**). Scale bar, 250 μm. The experiment was independently repeated 3 times with similar results for each mouse strain. **c,** Representative images showing dense distribution of mGFP and mRuby positive axons from the ZIr^Ecel1^ neurons to the PAG and from ZIm^Pde11a^ neuron to the PRF. Scale bar, 100 μm. **d,** Quantification of integrated fluorescence intensity of mRuby and mGFP-labeled axon terminals in PAG or PRF of ZIr^Ecel1^ and ZIm^Pde11a^ neurons. \**P* = 0.015, \*\**P* = 0.0032, n = 3 mice. **e,** Quantification of relative fluorescent density of mRuby and mGFP-labeled axon terminals of ZIr^Ecel1^ and ZIm^Pde11a^ neurons in the target brain regions. N = 9 slices from 3 mice for each brain region. Abbreviations: LS, lateral septal nucleus; POA, preoptic area; PVT, paraventricular thalamic nucleus; Re, reuniens thalamic nucleus; LH, lateral hypothalamic area; MD, mediodorsal thalamic nucleus; Po, posterior thalamic nucleus; Auv, secondary auditory cortex; APTD, anterior pretectal nucleus; Dk, nucleus of Darkschewitsch; RN, red nucleus; PAG, periaqueductal gray; PRF, pontine reticular nucleus, oral part; SC, superior colliculus. Two-tailed unpaired t test for (**d**). Data are presented as mean ± SEM. Source data are provided as a Source Data file.

### Activation of ZIr^Ecel1^ neurons and the GABAergic projection from ZIr^Ecel1^ to PAG pathway mediates self-grooming

To investigate the behavioral function of ZIr^Ecel1^ GABAergic projection neurons, we bilaterally injected AAV virus expressing Cre-dependent channelrhodopsin2-EYFP (AAV-DIO-ChR2-EYFP) or EGFP as control (AAV-DIO-EGFP) into the ZIr of Ecel1-Cre mice (Fig. 5a and Supplementary Fig. 4a). We found optogenetic activation of ChR2-expressing ZIr^Ecel1^ GABAergic neurons at different stimulation frequencies (5Hz, 10Hz, 20Hz) elicited robust self-grooming behavior compared with EGFP-expressing control mice (Fig. 5b-d, Supplementary Fig. 4b,c and Supplementary Video 1 and 2). High frequency optogenetic stimulation (20 Hz) also increased food intake or gnawing behavior (Supplementary Fig. 5a-d and Supplementary Video 3-6), consistent with previous findings on optogenetic activation of GABAergic neurons in the rostral ZI^34^. Self-grooming is reported to be associated with stress^52, 53^, increased anxiety^54–56^ or reward behaviors^57, 58^. We further measured the Ecel1-Cre mice in real time place preference test (RTPP) and found that optogenetic activation of ZIr^Ecel1^ neurons resulted in aversion to the optogenetic stimulation paired side (Supplementary Fig. 6e-g). These results indicate that ZIr^Ecel1^ neurons mediate self-grooming behaviors could be associated with negative emotion.

**Fig. 5:**
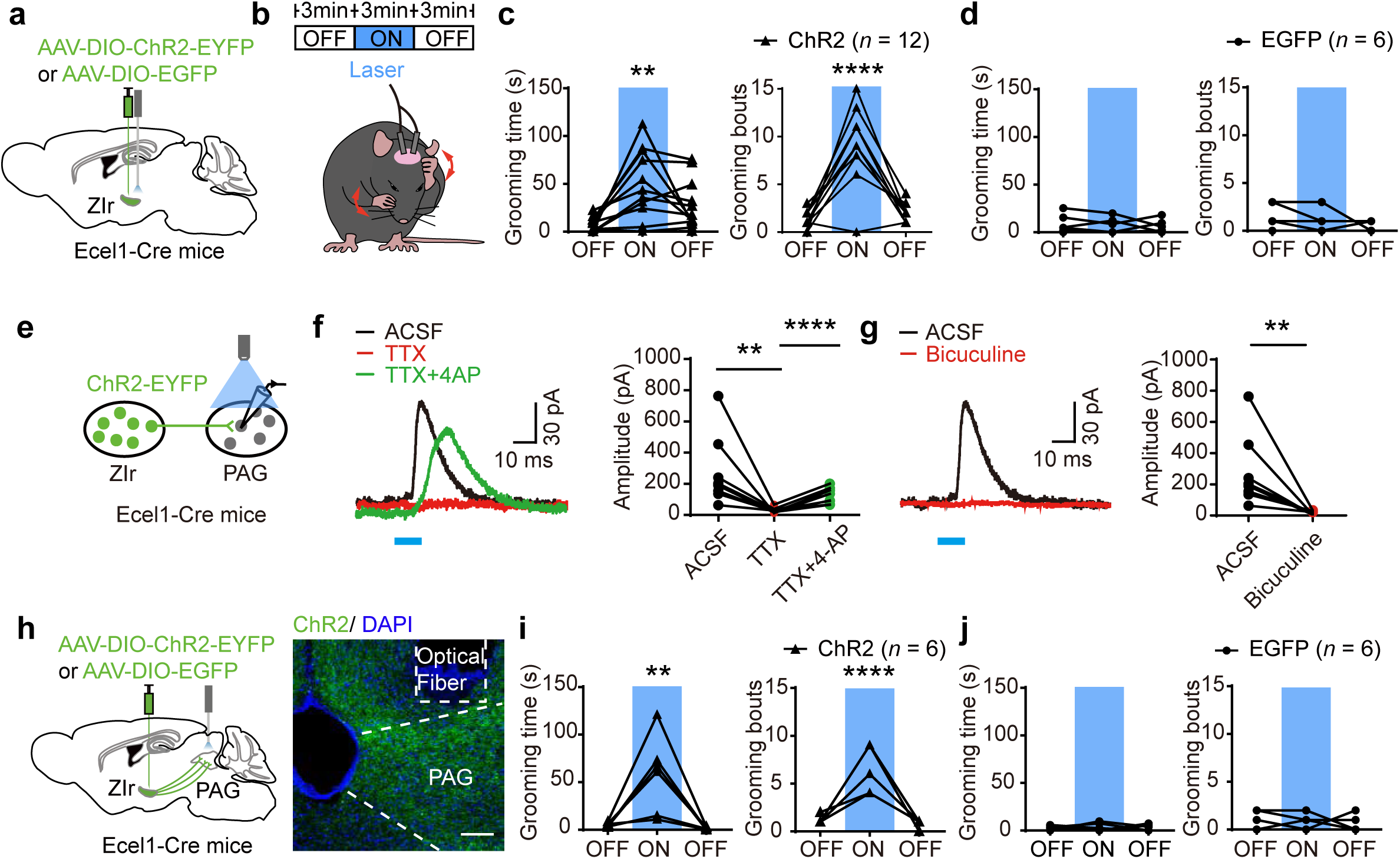
Optogenetic activation of ZIr^Ecel1^ neurons and the ZIr^Ecel1^ to PAG pathway both induce self-grooming. **a,** Schematic diagram showing optogenetic stimulation of the ChR2-EYFP or EGFP expressing ZIr^Ecel1^ neurons in Ecel1-Cre mice. **b,** Schematic diagram showing the behavioral paradigm for measurement of self-grooming behavior upon optogenetic stimulation. **c,** Statistic analysis of time spent for self-grooming (left) and grooming bouts (right) in 3 min before, during and after optogenetic stimulation of ZIr^Ecel1^ neurons in ChR2-group (**c**) and EGFP-group (**d**), respectively. Data are presented as mean ± SEM. \*\**P* = 0.0014, \*\*\*\**P* < 0.0001. Blue column indicates laser stimulation period (5 Hz, 3 min). **e,** Schematic diagram showing the recording of inhibitory postsynaptic currents (IPSCs) from PAG neurons evoked by optogenetic stimulation of ChR2-EYFP expressing axon terminals from the ZIr^Ecel1^ to the PAG in acute brain slices. **f,** Representative traces and statistical analysis of light-evoked IPSCs recorded in PAG neuron in ACSF and following sequential application of TTX (**f,** 1 μM. \*\**P* = 0.0082), TTX + 4-AP (**f,** 100 μM. \*\*\*\**P* < 0.0001), or bicuculline (**g,** 20 μM. \*\**P* = 0.0039). N = 9 neurons from 5 mice, two-tailed paired *t* test. Blue bar indicates optogenetic stimulation (10 ms). **h,** Schematic diagram and representative images showing optogenetic stimulation of the ChR2-EYFP or EGFP expressing terminals from ZIr^Ecel1^ to PAG pathway in Ecel1-Cre mice. Scale bar, 100 μm. **i,** Statistic analysis showing time spent for self-grooming (left) and grooming bouts (right) in 3 min before, during or after optogenetic stimulation of ZIr^Ecel1^ to PAG pathway in ChR2-group (**i**) and EGFP-group (**j**), respectively. \*\**P* = 0.0013, \*\*\*\**P* < 0.0001. Blue column indicates laser stimulation period (20 Hz, 3min). One-way ANOVA for (**c**), (**d**), (**i**) and (**j**). Two-tailed paired *t* test for (**f**) and (**g**). Data are presented as mean ± SEM. Source data are provided as a Source Data file.

Next, we examined the downstream brain region of ZIr^Ecel1^ neurons that mediated self-grooming behavior. Our results showed the axonal projection terminals in periaqueductal gray (PAG) from ZIr^Ecel1^ GABAergic neurons (Fig. 4c). The PAG neurons are reported to be an important sensory processing area which also involved in the regulation of self-grooming^54, 58^, different behaviors could be mediated by distinct neuron subtypes or ensembles of neurons driven by different descending afferents. To investigate whether the ZIr^Ecel1^ to PAG pathway mediate self-grooming, we bilaterally injected AAV-DIO-ChR2-EYFP or AAV-DIO-EGFP into the ZIr of Ecel1-Cre mice (Fig. 5h and Supplementary Fig. 7). We first examined the functional synaptic connection from the ZIr^Ecel1^ to PAG pathway by whole-cell patch-clamp recording of PAG neurons (Fig. 5e). We found that optogenetic stimulation of axon terminals from ZIr^Ecel1^ GABAergic neurons induced evoked inhibitory postsynaptic currents (IPSCs) in the PAG neurons in acute brain slices, that could be completely blocked by the GABA_A_ receptor antagonist bicuculline (20 μM). Administration of the Na^+^ channel inhibitor tetrodotoxin (TTX, 1 μM) blocked the optogenetically evoked IPSCs, while subsequent application of the K^+^ channel blocker 4-aminopyridine (4-AP, 100 μM) restored the evoked IPSCs (Fig. 5f, g). These results demonstrated functional monosynaptic GABAergic synaptic inputs from ZIr^Ecel1^ GABAergic neurons to the PAG.

In addition, we investigated the behavioral function of the ZIr^Ecel1^ to PAG pathway. We found optogenetic activation of the ChR2-expressing axon terminals from ZIr^Ecel1^ neurons to the PAG significantly induced increase in self-grooming time and grooming bouts (Fig. 5i, j). We also examined whether optogenetic stimulation of axonal projections from ZIr^Ecel1^ neurons to other target brain regions could induce self-grooming behavior. We found that optogenetic activation of the ChR2-expressing axon terminals from ZIr^Ecel1^ neurons to other targets including LS, POA, LH or SC had no discernible impact on either self-grooming time or self-grooming bouts in ChR2-group compared with EGFP-group (Supplementary Fig. 8). Together, these results demonstrate that activation of the ZIr^Ecel1^ neurons and the GABAergic projection from ZIr^Ecel1^ to PAG pathway mediates self-grooming behavior.

### ZIr^Ecel1^ neurons bidirectionally regulate self-grooming behavior

To monitor the activity of ZIr^Ecel1^ GABAergic neurons during self-grooming *in vivo,* we unilaterally injected AAV virus expressing Cre-dependent GCaMP6s and mCherry (as control) in the ZIr of Ecel1-Cre mice and recorded the calcium activity by fiber photometry (Fig. 6a, b). The self-grooming behavior can be efficiently induced by water spray toward the face^52, 58, 59^, we found the grooming time increased significantly compared to that before the water spray treatment (Fig. 6c, d). The calcium fluorescent signal of ZIr^Ecel1^ neurons increased before the start of grooming but decreased throughout the entire self-grooming period, and then returned to baseline after termination of self-grooming during both water spray induced and spontaneous self-grooming (Fig. 6e-g and Supplementary Fig. 9a-f). These results suggest that the increased activity of ZIr^Ecel1^ GABAergic neurons preceded the self-grooming behavior^53, 60^.

**Fig. 6:**
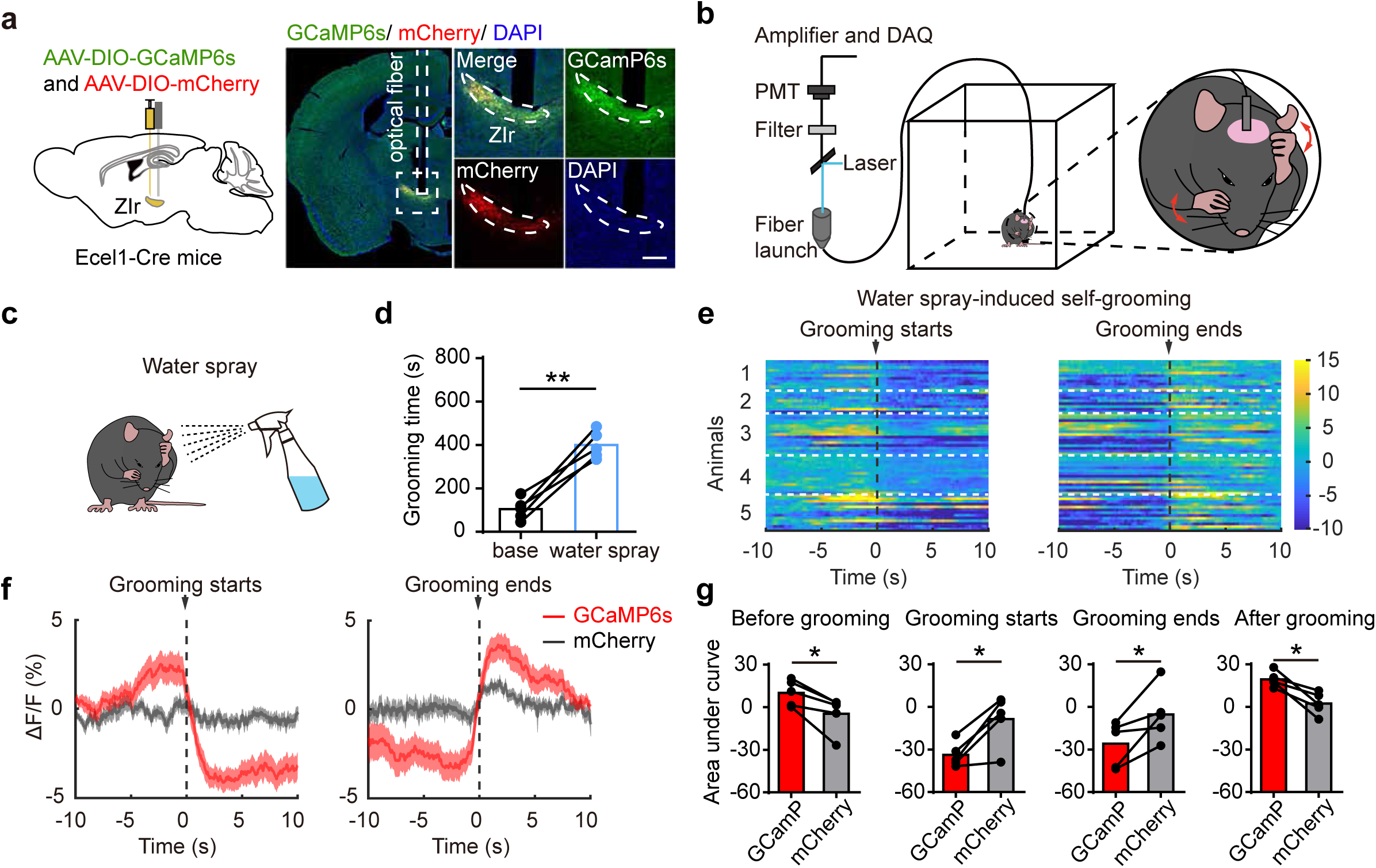
Calcium response patterns of Ecel1 positive neurons in the rostral ZI in self-grooming behavior. **a,** Schematic (left) and representative images showing GCamP6s and mCherry expression in ZIr of Ecel1-Cre mice (right). Scale bars, 500 μm and 250 μm (zoom-in image). The experiment was independently repeated 5 times with similar results. **b,** Schematic diagram showing the behavioral paradigm for fiber photometry recording in freely moving mice. **c,** Schematic diagram (**c**) and statistic analysis (**d**) showing water spray toward the face effectively induced self-grooming behaviors. \*\**P* = 0.0013, n = 5 mice. **e**, Heatmap (**e**) and average calcium transients (**f**) of ZIr^Ecel1^ neurons before, during and after water-spray induced self-grooming. Red trace: recording of GCaMP6s fluorescent signal; Gray trace: recording of control mCherry fluorescent signal. The dashed lines indicate the start or the end of self-grooming. Shaded areas around means indicate standard error of mean (SEM). **g,** Statistic analysis showing area under curve (AUC) of average calcium transients of GCamP6s channel compared with mCherry control channel before grooming starts (\**P* = 0.0156) and after grooming starts (\**P* = 0.0152), before grooming ends (\**P* = 0.0385) and after grooming ends (\**P* = 0.0114) in water-spray induced self-grooming. N = 5 mice per group. Two-tailed paired *t*-test for (**d**) and (**g**). Data are presented as mean ± SEM. Source data are provided as a Source Data file.

We further chemogenetically activated or inhibited the ZIr^Ecel1^ neurons by bilaterally injecting AAV-DIO-hM3Dq-mCherry or AAV-DIO-hM4Di-mCherry (or AAV-DIO-mCherry as control) into the ZIr of Ecel1-Cre mice (Fig. 7a, e). We found that chemogenetic activation of ZIr^Ecel1^ neurons significantly increased the self-grooming time and self-grooming bouts in hM3Dq-expressing mice compared with the mCherry-expressing control mice (Fig. 7b-d and Supplementary Fig. 9g), but chemogenetic inhibition of ZIr^Ecel1^ neurons showed no discernable effect on the spontaneous self-grooming behavior in hM4Di-expressing mice compared with the mCherry-expressing control mice (Supplementary Fig. 9h and i). To test whether ZIr^Ecel1^ neurons are required for water spray induced self-grooming behavior, the mice were i.p. administered with CNO (3 mg/kg) 30 min before water spray toward the mice face following by 20min self-grooming test (Fig. 7f). We demonstrated that chemogenetic inhibition of Ecel1 positive neurons in the ZIr significantly decreased the total self-grooming time and self-grooming bout duration in hM4Di-expressing mice compared with the mCherry-expressing control mice in water spray induced self-grooming (Fig. 7g-i and Supplementary Fig. 9k). Moreover, we further ablated the ZIr^Ecel1^ neurons by bilaterally injected AAV virus carrying Cre-dependent DTA (AAV-DIO-DTA) or mCherry (AAV-DIO-mCherry) as control into the ZIr of Ecel1-Cre mice (Fig. 7j and Supplementary Fig. 9l-n). While ablation of ZIr^Ecel1^ neurons showed no discernable effect on the spontaneous self-grooming behavior (Supplementary Fig. 9j), we found that ablation of ZIr^Ecel1^ neurons also significantly reduced the self-grooming time and grooming bouts induced by water spray in DTA-expressing mice compared with the mCherry-expressing control mice (Fig. 7k-m). These results indicates that the ZIr^Ecel1^ neurons can bidirectionally regulate the self-grooming behavior.

**Fig. 7:**
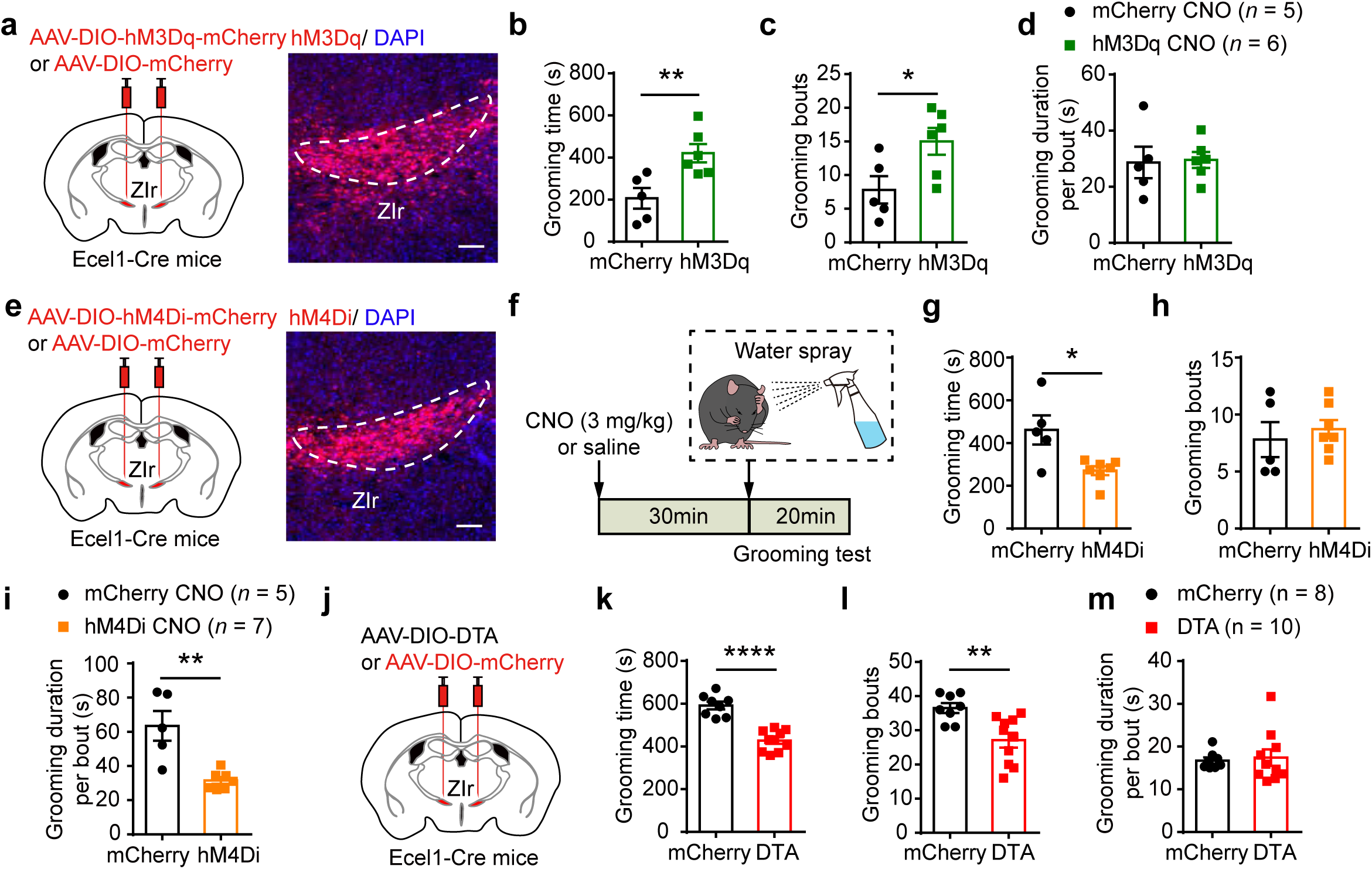
Ecel1 positive neurons in the rostral ZI are required for water spray induced self-grooming. **a,** Schematic diagram and representative images showing the stereotaxic bilateral injection of AAV-DIO-hM3Dq-mCherry or AAV-DIO-mCherry in the ZIr of Ecel1-Cre mice. Scale bar, 100 μm. The experiment was independently repeated 5 times with similar results. **b-d,** Statistic analysis of time spent for self-grooming (**b**), grooming bouts (**c**) and grooming duration per bout (**d**) in 20min following injection of CNO in hM3Dq-group mice compared with mCherry-group mice. \*\**P* = 0.0097, \**P* = 0.0336. **e,** Schematic diagram and representative images showing the injection of AAV-DIO-hM4Di-mCherry to chemogenetically inhibit ZIr^Ecel1^ neurons of Ecel1-Cre mice. Scale bar, 100 μm. The experiment was independently repeated 7 times with similar results. **f,** The behavioral paradigm for measurement of water-spray induced self-grooming behavior following injection of CNO in hM4Di-group or mCherry-group mice. **g-i,** Statistic analysis of time spent for water-spray induced self-grooming (**g**), grooming bouts (**h**) and grooming duration per bout (**i**) in 20 min following injection of CNO in hM4Di mice. \**P* = 0.0117, \*\**P* = 0.0018. **j,** Schematic diagram showing the stereotaxic bilateral injection of AAV-DIO-DTA to ablate the ZIr^Ecel1^ neurons of Ecel1-Cre mice. **k-m,** Statistic analysis of water-spray induced self-grooming time (**k**), grooming bouts (**l**) and grooming duration per bout (**m**) in 20 min following water spray in DTA-group mice. \*\*\*\**P* < 0.0001, \*\**P* = 0.0036. Two-tailed unpaired *t*-test for (**b-d**), (**g-i**) and (**k-m**). Data are presented as mean ± SEM. Source data are provided as a Source Data file.

### Activation of ZIm^Pde11a^ neurons and the GABAergic projection from ZIm^Pde11a^ to PRF pathway promotes transition from sleep to wakefulness

We next investigated the behavioral function of ZIm^Pde11a^ GABAergic neurons by injecting AAV-DIO-ChR2-EYFP or control AAV-DIO-EGFP virus into the ZIm of Pde11a-Cre mice (Supplementary Fig. 10a). Optogenetic activation of ZIm^Pde11a^ neurons during the awake state of mice did not have significant impact on the center time or locomotor activity in open field test, or the time spent in open arm in EPM test, or preference to light stimulation in RTPP test (Supplementary Fig. 10f-l). Our result showed extensive axonal projections from the ZIm^Pde11a^ neurons to the oral part of pontine reticular formation (PRF) (Fig. 4c), which is a component of the ascending reticular activating system that modulates rapid eye movement sleep (REM) in sleep and wakefulness cycle^61, 62^. To investigate whether ZIm^Pde11a^ GABAergic projection neurons participated in regulating sleep-wakefulness cycle, we monitored the activity of ZIm^Pde11a^ neurons by AAV mediated Cre-dependent expression of GCaMP6s or mCherry (as control) using fiber photometry. The phase of sleep-wakefulness cycle was determined by electroencephalogram (EEG) and electromyogram (EMG) recordings (Fig. 8a, b). We found the ZIm^Pde11a^ neurons are active during REM sleep and wakefulness, the calcium signal increases during the transition from NREM to REM sleep and decreases during the transition from REM sleep to wakefulness and wakefulness to NREM sleep (Fig. 8c-e).

**Fig. 8:**
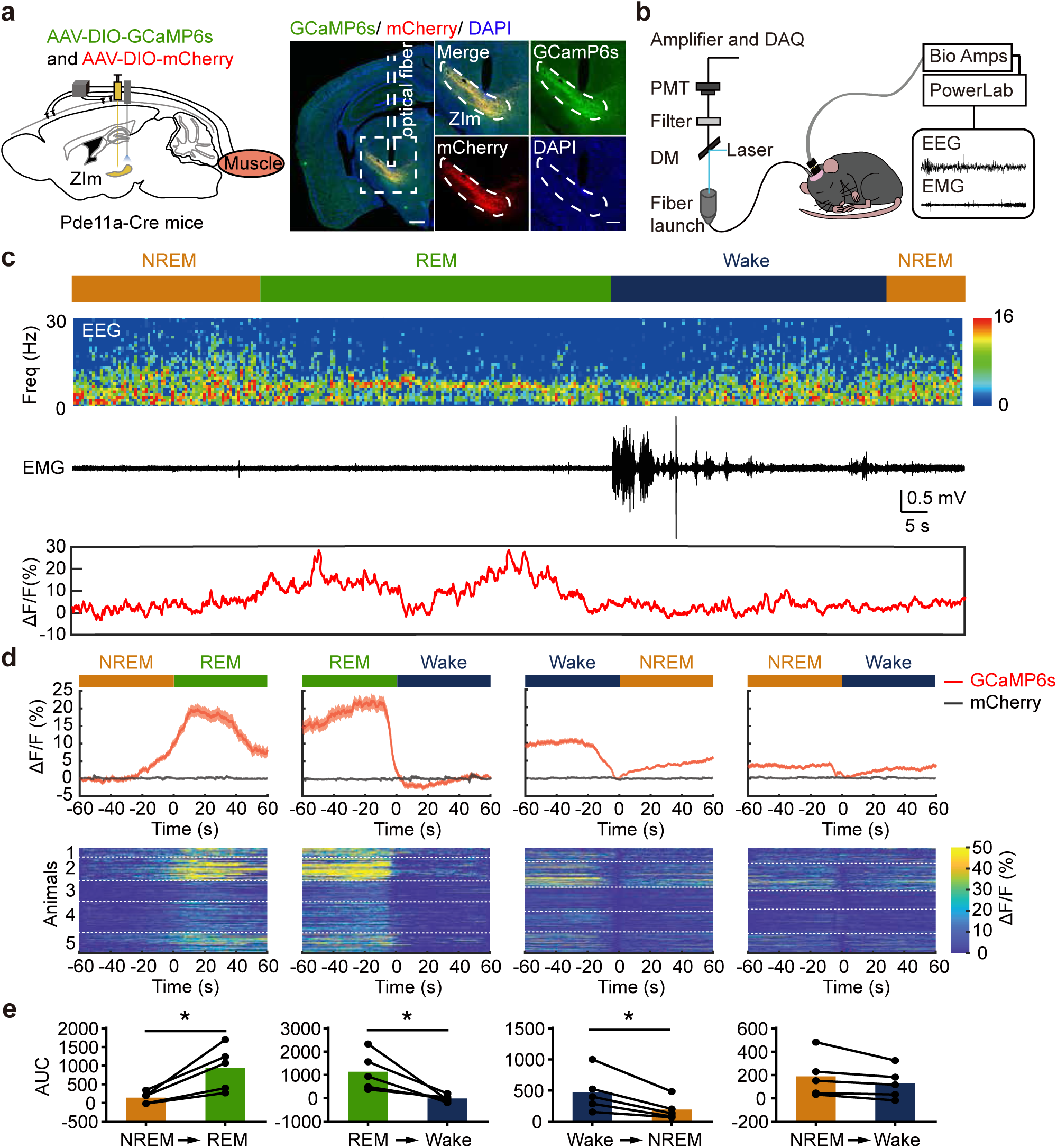
Calcium response patterns of Pde11a positive neurons in the medial ZI during sleep-wakefulness cycle. **a,** Schematic and representative images (right) showing *in vivo* recording of calcium response pattern of GCamP6s- or mCherry-expressing Pde11a positive neurons in ZIm. Gray column indicates the position of optical fiber. Scale bars, 500 μm and 250 μm (zoom-in image). The experiment was independently repeated 5 times with similar results. **b,** Schematic diagram showing the behavioral paradigm for fiber photometry, electroencephalogram (EEG) and electromyographic (EMG) recordings in freely moving mice. **c,** Sample trace showing the changes of fluorescent calcium signal of ZIm^Pde11a^ neurons aligned to sleep-wakefulness state transitions. NREM (yellow bar), non-rapid eye movement; REM (green bar), rapid eye movement; Wake (blue bar). **d,** Average trace and heatmap showing changes of the fluorescent calcium signal of ZIm^Pde11a^ neurons during sleep-wakefulness state transitions. The red trace showed average fluorescent GCaMP6s or control mCherry (gray) signals, while shaded areas indicate standard error of mean (SEM). **e,** Statistical analysis of the area under curve (AUC) of average fluorescent calcium signal of ZIm^Pde11a^ neurons during sleep-wakefulness state transitions. N = 5 mice per group. \**P* = 0.0207 (NREM to REM), \**P* = 0.0332 (REM to Wake), \**P* = 0.0171 (Wake to NREM), two-tailed paired *t*-test. Data are presented as mean ± SEM. Source data are provided as a Source Data file.

Optogenetic activation of ChR2-expressing ZIm^Pde11a^ GABAergic projection neurons promotes the transition from NREM or REM sleep to wakefulness in comparison with the EGFP control group (20 Hz, 120 seconds per trial, applied every 10–15 minutes. Fig. 9a-d and Supplementary Video 7 and 8). The transition probability from REM sleep to wakefulness and NREM sleep to wakefulness was significantly increased upon optogenetic activation of the ZIm^Pde11a^ GABAergic projection neurons, along with a complementary decrease of transition probability from wakefulness to NREM or NREM to REM sleep (Fig. 9e and Supplementary Fig. 10b-e). We verified the functional synaptic connectivity from ZIm^Pde11a^ to PRF pathway by whole-cell patch-clamp recording of neurons in the PRF (Fig. 9f). Optogenetic stimulation of the ChR2-expressing axonal terminals from ZIm^Pde11a^ neurons evoked IPSCs in PRF neurons that can be blocked by bicuculline (20 μM) or TTX (1 μM). The blockade of IPSCs by TTX was restored following application of 4-AP (100 μM), suggesting monosynaptic GABAergic inputs from ZIm^Pde11a^ to PRF pathway (Fig. 9g, h). We investigated the behavioral function of the ZIm^Pde11a^ to PRF pathway and found optogenetic activation of the GABAergic projections from ZIm^Pde11a^ to the PRF increased the transition probability from NREM to wakefulness and decreased the transition probability from wakefulness to NREM or from NREM to REM sleep (Fig. 9i-l and Supplementary Fig. 11). Thus, these results demonstrate that activation of the ZIm^Pde11a^ GABAergic projection neurons and the ZIm^Pde11a^ to PRF pathway promotes transition from sleep to wakefulness.

**Fig. 9:**
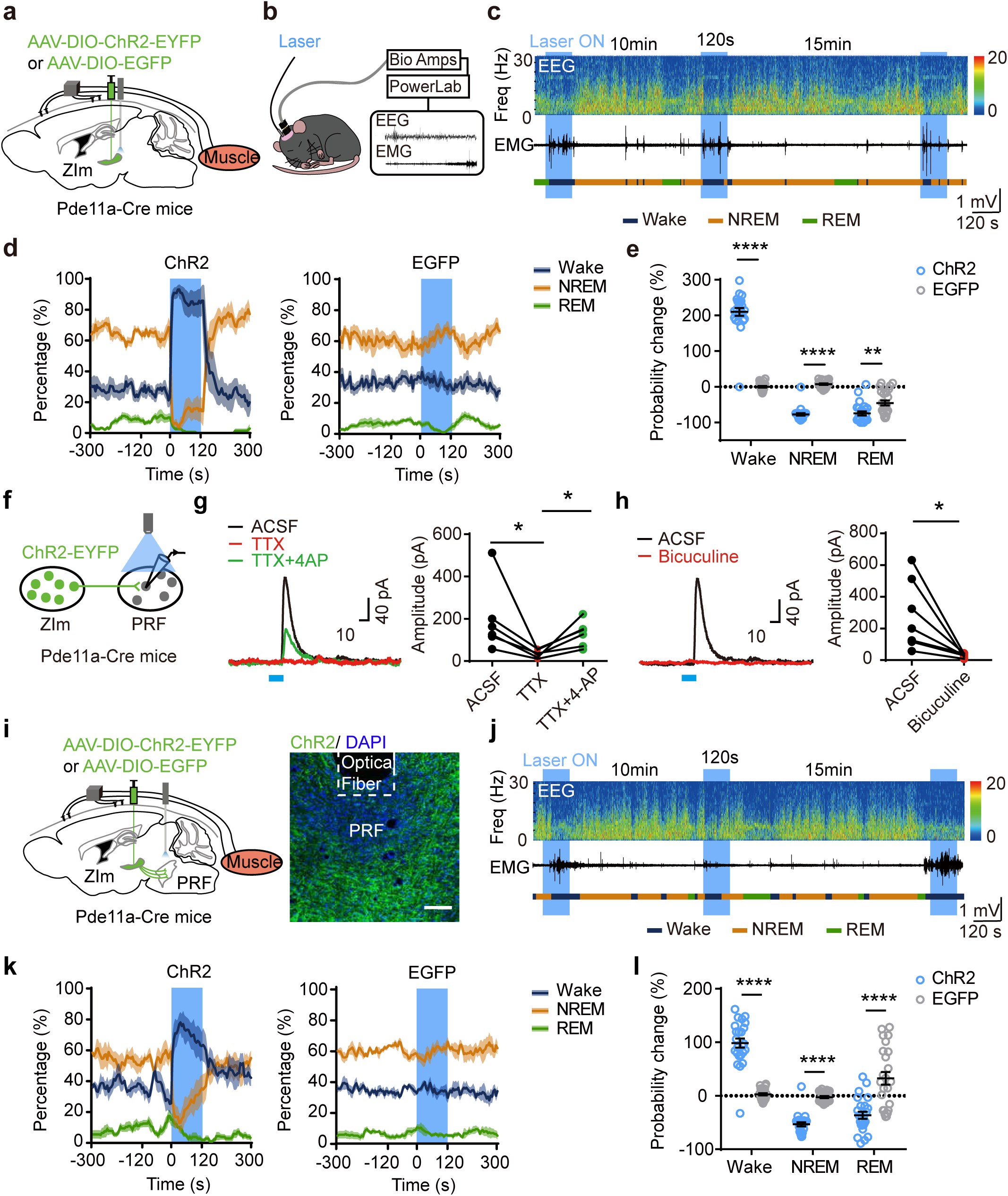
Optogenetic activation of ZIm^Pde11a^ neurons promotes wakefulness through the ZIm^Pde11a^ to PRF pathway. **a,** Optogenetic stimulation of ChR2-EYFP or EGFP expressing ZIm^Pde11a^ neurons and the behavioral paradigm for EEG-EMG recordings of freely moving Pde11a-Cre mice (**b**). **c,** Sample EEG power spectrum, EMG traces and brain states (color coded) during optogenetic activation of ZIm^Pde11a^ neurons. Dark blue: wake; Yellow: NREM; Green: REM. Freq: frequency. **d,** The percentage of wake, NREM, or REM states before, during, and after optogenetic stimulation of ZIm^Pde11a^ neurons in ChR2-group (left, n = 5 mice) and EGFP-group (right, n = 6 mice). Shaded areas around means indicate standard error of mean (SEM). **e,** Probability change of each state 120s before and during optogenetic stimulation of ZIm^Pde11a^ neurons in ChR2-group compared with EGFP-group. \*\*\*\**P* < 0.0001, \*\**P* = 0.0021. **f-h,** Schematic diagram (**f**) and representative traces (**g** and **h**, left) showing the recording of evoked IPSCs in PRF neurons in acute brain slices. TTX, \**P* = 0.0146; TTX+4-AP, \**P* = 0.0129 (n = 6 neurons from 4 mice); Bicuculline, \**P* = 0.0215 (n = 7 neurons from 4 mice). Blue bar indicates laser stimulation period (10ms). **i,** Optogenetic stimulation of ChR2-EYFP or EGFP expressing terminals of ZIm^Pde11a^ to PRF pathway together with EEG-EMG recordings in Pde11a-Cre mice. Scale bar, 100μm. The experiment was independently repeated 5 times with similar results. **j,** Sample EEG power spectrum, EMG traces and brain states during optogenetic activation of ZIm^Pde11a^ to PRF pathway. **k,** The percentage of wake, NREM, or REM states before, during, and after optogenetic stimulation of ZIm^Pde11a^ to PRF pathway in ChR2-group (left, n = 5 mice) and EGFP-group (right, n = 5 mice). Shaded areas around means indicate standard error of mean (SEM). **l,** Probability change of each state 120s before and during optogenetic stimulation of ZIm^Pde11a^ to PRF pathway in ChR2-group compared with EGFP-group. \*\*\*\**P* < 0.0001. Two-tailed unpaired t test for (**e**) and (**l**). Two-tailed paired t test for (**g**) and (**h**). Data are presented as mean±SEM. Blue bar indicates laser stimulation period (20Hz, 120s). Source data are provided as a Source Data file.

### ZIm^Pde11a^ neurons bidirectionally regulate sleep-wakefulness cycle

We further chemogenetically activated ZIm^Pde11a^ neurons by bilaterally injecting AAV-DIO-hM3Dq-mCherry or AAV-DIO-mCherry as control into the ZIm of Pde11a-Cre mice (Fig. 10a). We found that chemogenetic activation of ZIm^Pde11a^ neurons significantly increased wakefulness and decreased NREM and REM sleep in hM3Dq-expressing mice following CNO injection compared with saline injection group during the light phase (Fig. 10b-h). In addition, the number of episodes during wakefulness and NREM both increases, but the duration of average NREM episode significantly decreases, suggesting chemogenetic activation of ZIm^Pde11a^ neurons increased wakefulness at the expense of sleep time and sleep quality in the light phase (Fig. 10i, j). Chemogenetic inhibition did not alter the sleep-wakefulness state in hM4Di-expressing mice or in mCherry-expressing control mice (Supplementary Fig. 12a-h). To determine whether ZIm^Pde11a^ neurons are required for sleep-wakefulness transition, we selectively ablated the ZIm^Pde11a^ neurons by bilaterally injected AAV-DIO-DTA or AAV-DIO-mCherry as control into the ZIm of Pde11a-Cre mice (Fig. 10k) and verified the efficiency of DTA ablation (Supplementary Fig. 12i-k). We found that ablation of ZIm^Pde11a^ neurons significantly decreased wakefulness and increased the NREM and REM sleep during dark phase (Fig. 10l-q). In addition, the number of episodes during wakefulness, NREM and REM all increases, but the duration of average wakefulness episode significantly decreases, suggesting ZIm^Pde11a^ neurons may be required for wakefulness in the dark phase (Fig. 10r, s). These results demonstrate that the ZIm^Pde11a^ neurons can bidirectionally regulate the sleep and wakefulness cycle.

**Fig. 10:**
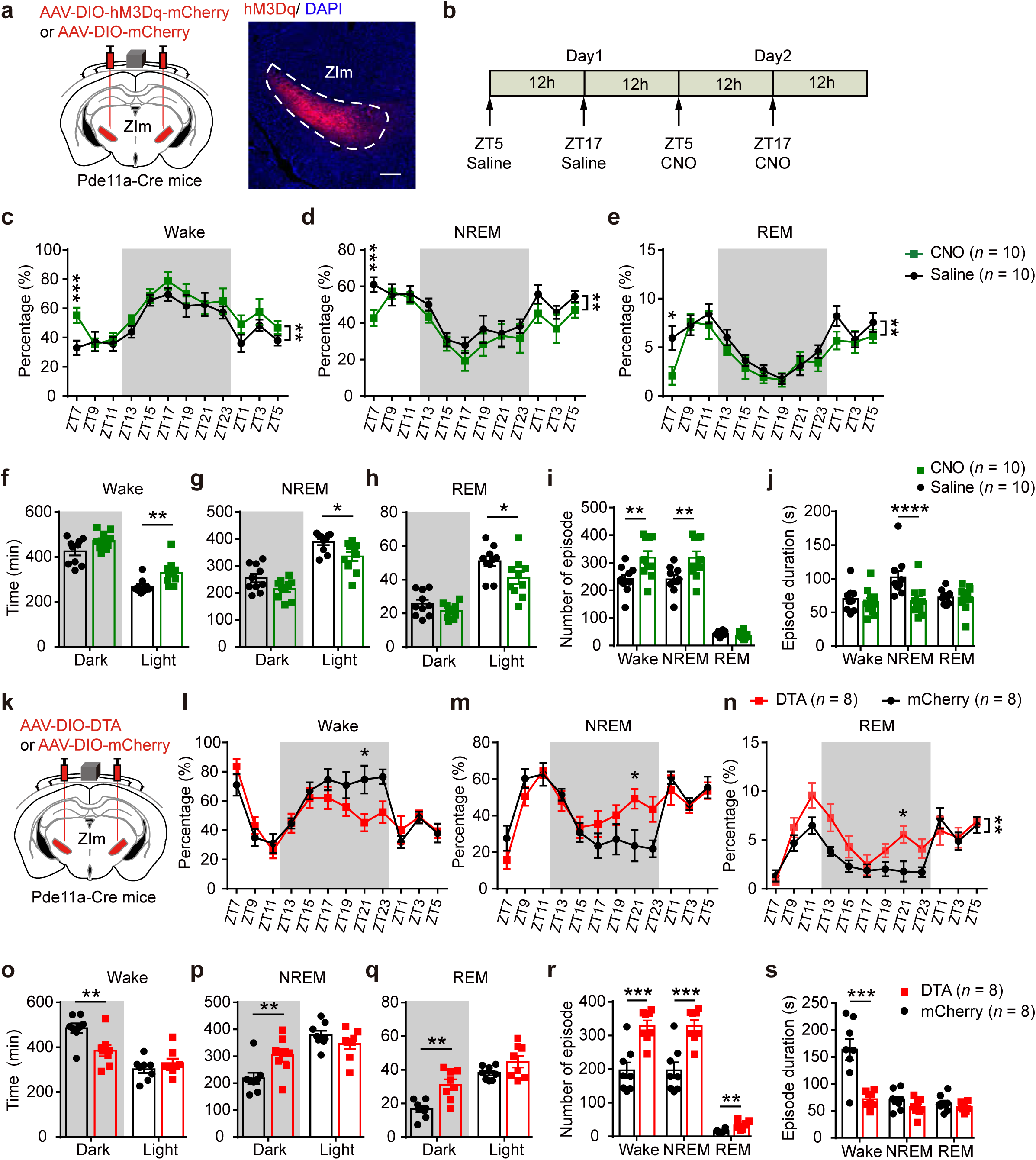
Selective chemogenetic activation or ablation of ZIm^Pde11a^ neurons bidirectionally regulate sleep-wakefulness cycle. **a,** Schematic diagram and representative images showing the injection of AAV-DIO-hM3Dq-mCherry or AAV-DIO-mCherry into the ZIm of Pde11a-Cre mice. Scale bar, 250μm. The experiment was independently repeated 10 times with similar results. **b,** Schematic showing sleep recording conditions with saline- or CNO-injection in hM3Dq-group mice. **c-e,** The percentage of time spent in wake (**c,** ZT7, \*\*\**P*=0.0008), NREM sleep (**d**, ZT7, \*\*\**P*=0.0007) and REM sleep (**e**, ZT7, \**P*=0.0117) in 2-h bins following injection of saline or CNO in hM3Dq-group mice. Wake, \*\**P*=0.0056 (**c**); NREM, \*\**P*=0.0080 (**d**); REM, \*\**P*=0.009 (**e**). **f-h,** Time spent in wake (**f**), NREM sleep (**g**) and REM sleep (**h**) across the 24-hour sleep-wake cycle in hM3Dq-group mice. Wake, \*\**P*=0.0083 (**f**); NREM, \**P*=0.0180 (**g**); REM, \**P*=0.0304 (**h**). **i,** Number and duration (**j**) of wake, NREM sleep and REM sleep episodes during the light phase. Number: Wake, \*\**P*=0.0027; NREM, \*\**P*=0.0026. Duration: NREM, \*\*\*\**P*<0.0001. **k,** Schematic diagram showing the injection of AAV-DIO-DTA to ablate the ZIm^Pde11a^ neurons. **l-n,** The percentage of time spent in wake (**l**, ZT21, \**P*=0.0165), NREM sleep (**m**, ZT21, \**P*=0.0244) and REM sleep (**n**, ZT21, \**P*=0.0427) in 2-h bins in DTA-group mice compared with mCherry-group mice. REM, \*\**P*=0.0040 (**n**). **o-q,** Time spent in wake (**o**), NREM sleep (**p**) and REM sleep (**q**) across the 24-hour sleep-wake cycle in DTA-group mice compared with mCherry-group mice. Wake, \*\**P*=0.0045 (**o**); NREM, \*\**P*=0.0082 (**p**); REM, \*\**P*=0.0013 (**q**). **r,** Number and duration (**s**) of wake, NREM sleep and REM sleep episodes during the dark phase. Number: Wake, \*\*\**P*=0.0004; NREM, \*\*\**P*=0.0005; REM, \*\**P*=0.0016. Duration: Wake, \*\*\**P*=0.0004. Two-way ANOVA and Sidak’s multiple comparisons test for (**c-e**) and (**l-n**). Two-tailed paired t test for (**c-j**). Two-tailed unpaired t test for (**o-s**). Data are presented as mean±SEM. Grey bar indicates the dark phase (ZT12-ZT0). Source data are provided as a Source Data file.

We further explored whether ZIr^Ecel1^ and ZIm^Pde11a^ neurons play distinct roles in regulating self-grooming or sleep-wakefulness cycle behaviors. We found that ablation of the ZIm^Pde11a^ neurons by selective expression of DTA had no effects on the spontaneous or water spray induced self-grooming behaviors compared with mCherry control group (Supplementary Fig. 13j-m). Additionally, ablation of ZIr^Ecel1^ neurons by DTA had no significant impact on the time or episodes of wakefulness, NREM sleep or REM sleep compared with mCherry-group mice (Supplementary Fig. 13a-i). These results suggest that these ZIr^Ecel1^ and ZIm^Pde11a^ neurons differentially regulate self-grooming or sleep-wakefulness cycle behaviors, respectively.

Taken together, our study revealed the molecular markers and spatial distribution of GABAergic neuron subtypes in subregions of zona incerta. By *in vivo* circuit mapping and manipulating the neuronal activity, we further delineated distinct functions for two discrete GABAergic projection neuron subtypes: the ZIr^Ecel1^ GABAergic neurons regulate self-grooming through projection to PAG pathway and ZIm^Pde11a^ GABAergic neurons promote transition from sleep to wakefulness through projection to PRF pathway. These results demonstrated the molecular markers, spatial distribution and circuit mechanisms of GABAergic projection neuron subtypes in the ZI that mediate distinct innate behaviors.

## Discussion

### Molecular markers and spatial localization of GABAergic projection neuron subtypes in zona incerta

Despite the unique long-range axonal projection pattern and distinct functions of GABAergic projection neurons, the lack of specific molecular markers makes it difficult to selectively target or to investigate the function of GABAergic projection neurons *in vivo*. By single-nucleus transcriptomic sequencing of the ZI and quantifying the spatial expression of marker genes of GABAergic neurons in different subregions along the anteroposterior axis of the ZI, we found two discrete *Ecel1-* and *Pde11a-*expressing GABAergic neurons that are non-overlapping and exhibit dominant expression in the rostral and medial ZI, respectively (Fig. 2d, e).

By *in vivo* circuit mapping, we further demonstrated that ZIr^Ecel1^ and ZIm^Pde11a^ neurons sent long-range GABAergic efferent projections to segregated distant brain regions in the forebrain and the midbrain (Fig. 4 and Supplementary Fig. 3c), implying distinct behavioral functions. We also found the axonal projections from the ZIm^Pde11a^ GABAergic projection neurons to the layer 1 of auditory cortex (Supplementary Fig. 3c), consistent with recent finding reporting the top-down GABAergic projection from ZI in regulating neocortical associative fear learning^27^. The establishment and maintenance of specific projections are determined by both genetic and environmental factors^63–65^. Recent studies showed heterogeneity of the projection area and potential functions in transcriptomic neuron subtypes^66^. Therefore, the same transcriptomic neuronal subtypes could establish different efferent projections during development^66^. The diverse behavioral functions of the ZI are routed in the heterogenous neuron populations, that are possibly generated from different progenitor cells during development^67, 68^. Future studies are warranted to elucidate the developmental origin of GABAergic neuron subtypes in the ZI.

### Activation of ZIr^Ecel1^ GABAergic projection neurons induced self-grooming behaviors may be associated with stress

Previous reports suggested that ZI neurons could receive information from different sensory modalities including auditory, whisker and visual stimulation^31, 37, 69^. By monitoring the calcium dynamics of ZIr^Ecel1^ neurons *in vivo*, we found increased calcium activity of ZIr^Ecel1^ neurons before water spray induced self-grooming, suggesting these neurons could be activated by water spray induced unpleasant sensory stress^52, 59^ (Fig. 6). Chemogenetic inhibition or ablation of ZIr^Ecel1^ neurons impaired the self-grooming behavior induced by water spray but had little effect on the spontaneous self-grooming behavior (Fig. 7e-m and Supplementary Fig. 9h-j). Thus, these results suggest that activation of ZIr^Ecel1^ neurons induced self-grooming behaviors could be associated with sensory stress.

Self-grooming is an innate behavior that could be elicited by comfort or stressful conditions^70^. Our results demonstrated that optogenetic activation of ZIr^Ecel1^ neurons increased the bouts of self-grooming and induced aversion to laser stimulation paired side (Fig. 5a-d and Supplementary Fig. 6e-g), resembling with stress-associated self-grooming behavior that often accompanied by negative emotional state^52, 59^. Previous study showed that activation of somatostatin (SST)-expressing neurons or calretinin (CR)-expressing neurons in the ZI differentially modulate anxiety^40^. We found that only subsets of *Ecel1*-expressing neurons co-expressed *SOM* (*Ecel1^+^SST^+^/Ecel1^+^*, 23.12 ± 1.12%; *Ecel1^+^SST^+^/SST^+^*, 33.93 ± 0.96%), while *Ecel1*-expressing neurons rarely co-expressed *CR* in the rostral ZI (*Ecel1^+^*CR^+^/*Ecel1^+^*, 4.10 ± 0.15%; *Ecel1^+^*CR^+^/CR^+^, 13.80 ± 2.38%. Supplementary Fig. 14). In addition, optogenetic stimulation of ZIr^Ecel1^ neurons did not affect the center time in open field behavioral test, or time spent in the open arm in elevated plus maze (EPM) behavioral test (Supplementary Fig. 6a-d), or the time of investigating the novel object compared with familiar object in novel object recognition test (Supplementary Fig. 6h, i). Together, these results suggest that ZIr^Ecel1^ neurons could be a distinct subpopulation of GABAergic projection neurons that mediated stress-related self-grooming behavior.

Stress may lead to decreased feeding and increased grooming^71, 72^. Previous studies demonstrated that hypothalamic neuronal circuit can exert scalable control over feeding and negative emotional states^52^. Our results showed that low frequency optogenetic stimulation (5Hz) of ZIr^Ecel1^ neurons only promote self-grooming, while high frequency optogenetic stimulation (20Hz) of ZIr^Ecel1^ neurons also increased feeding, suggesting ZIr^Ecel1^ neurons may orchestrate behaviors related to stress response and feeding (Fig. 5b-d, Supplementary Fig. 4b,c and Supplementary Fig. 5a-d). In addition, we found ablation of ZIr^Ecel1^ neurons significantly reduced water spray induced self-grooming (Fig. 7j-m), but had no discernible impact on food intake or the body weight (Supplementary Fig. 5e-h). These results demonstrate that ZIr^Ecel1^ neurons are required for sensory stress related self-grooming.

### ZIm^Pde11a^ GABAergic projection neurons participate in regulating sleep-wakefulness cycle

Extensive GABAergic projections from the diencephalon including ZI, reticular thalamus and hypothalamus to pontine reticular formation (PRF) have been identified by retrograde tracing approaches^73^. Endogenous GABA levels were increased during wakefulness and application of GABA_A_ receptor agonists in PRF promotes wakefulness^74–76^. We demonstrated that activation of ZIm^Pde11a^ neurons and the GABAergic projection from ZIm^Pde11a^ to PRF promote the transition from sleep to wakefulness (Fig. 9). We found increased Ca^2+^ activity of ZIm^Pde11a^ neurons during REM sleep and in wakefulness using fiber photometry recordings (Fig. 8). Recent studies using *in vivo* micro-endoscopic imaging showed that GABAergic neurons in the ZI demonstrated heterogenous activity in sleep-wakefulness cycle, including neurons showing maximal Ca^2+^ activity during REM sleep (REM-active), neurons showing maximal Ca^2+^ activity during wakefulness (Wake-active), or neurons showing Ca^2+^ activity in both REM sleep and wakefulness (Wake/REM-active)^77^.

Activation of REM-active neurons promote REM sleep, often at the expense of NREM sleep^78, 79^, while activation of Wake/REM-active neurons (arousal neurons) promote wakefulness^80–82^. The ZIm^Pde11a^ neurons may include both REM-active and Wake/REM-active neurons as demonstrated by fiber photometry recording (Fig. 8). Therefore, optogenetic stimulation of ZIm^Pde11a^ neurons promotes the transition from sleep to wakefulness could be mediated by the activation of the Wake/REM-active ZIm^Pde11a^ neurons. Additionally, optogenetic activation of ZIm^Pde11a^ neurons decreased the investigating time of both familiar and novel object compared with control group in the novel object recognition test, (Supplementary Fig. 10m,n), suggesting a potential shift to alert state during wakefulness^83, 84^.

The zona incerta has been a target for chronic deep brain stimulation in treating Parkinson’s disease (PD)^85, 86^. Previous studies reported activation of Lhx6-expressing GABAergic neurons subtypes induced transition from wakefulness to sleep^35^. Our results showed that *Pde11a*-positive neurons rarely co-expressed with *Lhx6* (*Pde11a^+^Lhx6^+^/Pde11a^+^*, 2.23 ± 1.18%; *Pde11a^+^Lhx6^+^/Lhx6^+^*, 1.38 ± 0.69%, n = 6 brain slices from 3 mice. Supplementary Fig. 15), suggesting ZIm^Pde11a^ is a distinct GABAergic projection neuron subpopulation that promote the transition from sleep to wakefulness. These findings demonstrate that *Pde11a^+^* and *Lhx6^+^* neuronal subpopulations in the ZIm may differentially regulate sleep-wakefulness cycle. Intriguingly, sleep abnormalities are reported in Parkinson’s patients and in rodent models of PD^87, 88^, it is plausible that chronic deep brain stimulation of ZI may help improve sleep-related symptoms in Parkinson’s disease. Future studies are warranted to further elucidate the subtype and the exact role of GABAergic neurons in regulating sleep disturbances.

In summary, our results revealed the spatial distribution of two discrete populations of GABAergic projection neurons in segregated subregions of the ZI. By constructing transgenic mice to selectively target the ZIr^Ecel1^ and ZIm^Pde11a^ GABAergic projection neurons, we further demonstrated their efferent axonal projection to non-overlapping distant brain regions and mediate distinct innate behavioral functions. These findings uncovered the molecular markers, spatial organization and functions for GABAergic projection neuron subtypes in the zona incerta that play distinct roles in regulating self-grooming and sleep-wakefulness cycle.

## Methods

### Animals

All experiments were conducted in accordance with the guidelines of Zhejiang University (ZJU) and were approved by the Animal Advisory Committee at ZJU. Wild-type C57BL/6J (JAX# Stock No. 000664) and transgenic GAD67-GFP mice (JAX# Stock No. 007677) were obtained from Jackson Laboratories. Transgenic mice Ecel1-Cre and Pde11a-Cre (C57/BL6 background) were generated by CRISPR/Cas9 technology at the Shanghai Model Organisms Center (China) or Biocytogen (Beijing) company. All mice were bred and housed at the temperature (23 ± 1 °C) and humidity (40%) controlled animal facility at ZJU under a 12 h light/dark cycle with light on at zeitgeber time 0 (ZT 0; 7:00), food and water were available ad libitum. Only adult (>8 weeks old) mice of normal appearance and weight were used for behavioral tests. Littermate mice were split into random groups before AAV viral injection.

### Viruses

AAV2/9-DIO-mGFP-Synaptophysin-mRuby (titer: 8.9×10^12^ genomic copies per ml), AAV2/9-EF1α-DIO-hChR2-EYFP (titer: 2×10^12^ genomic copies per ml) were purchased from Taitool Bioscience, China. AAV2/9-hSyn-DIO-EGFP (titer: 1.44×10^12^ genomic copies per ml) were purchased from WZ Bioscience, China. AAV2/9-EF1α-DIO-GCamP6s (titer: 8.35×10^12^ genomic copies per ml). AAV2/9-CAG-DIO-DTA (titer: 6.13×10^12^ genomic copies per ml). AAV2/9-DIO-mCherry (titer: 5.48×10^12^ genomic copies per ml) were purchased from BrainVTA Bioscience, China. AAV2/9-hSyn-DIO-hM3Dq-mCherry (titer: 3.27×10^13^ genomic copies per ml). AAV2/9-hSyn-DIO-hM4Di-mCherry (titer: 8.53×10^13^ genomic copies per ml) were purchased from Sunbio Medical Biotechnology Company, China.

### Stereotaxic injection

Adult mice were anesthetized with pentobarbital sodium solution (50 mg/kg) via an intraperitoneal (i.p.) injection. Mice were placed on a stereotaxic frame (RWD Life Science, 68046) and anesthesia were maintained with 1%-1.5% isoflurane during surgery. Injection of AAV virus was performed with a manual viral syringe pump connected to glass micropipette with a 10-50 μm diameter tip (Drummond Scientific Company, Wiretrol II). After each injection, the syringe was left *in situ* for an additional 10 min and then withdrawn slowly. Mice were randomly assigned to experimental and control groups before virus injection.

To map the axonal projection pattern of GABAergic neurons in the rostral or medial zona incerta, anterograde tracing virus AAV2/9-DIO-mGFP-Synaptophysin-mRuby (80 nl. nl: nano-liter) was unilaterally injected into the ZIr (anteroposterior (AP): −1.00 mm, mediolateral (ML): −0.60 mm, dorsoventral (DV): −4.45 mm) of Ecel1-Cre mice or the ZIm (AP: −2.30 mm, ML: −1.50 mm, DV: −4.20 mm) of Pde11a-Cre mice. Three weeks after injection, mice were sacrificed and brain sections were collected for fluorescent imaging. The projection pattern was shown as the percentage of total relative fluorescent density occupied by different downstream brain areas.

The neuronal activity of GABAergic neurons was monitored by in *vivo* fiber photometry recording. AAV2/9-EF1α-DIO-GCamP6s and AAV2/9-DIO-mCherry were injected as a mixture at 2:1 ratio (80 nl) into the ZIr of Ecel1-Cre mice or ZIm of Pde11a-Cre mice. Three weeks after injection, optical fibers (200 μm O.D., 0.37 NA, Inper, Hangzhou, China) were implanted above the ZIr in Ecel1-Cre mice or ZIm in Pde11a-Cre mice, respectively.

To activate GABAergic neuron subtypes by in *vivo* optogenetic stimulation, AAV2/9-EF1α-DIO-hChR2-EYFP or AAV2/9-hSyn-DIO-EGFP (80 nl) were bilaterally injected into the ZIr of Ecel1-Cre mice or unilaterally injected into the ZIm of Pde11a-Cre mice. Three weeks after injection, optical fibers were implanted at a 10° angle relative to the vertical plane above ZIr bilaterally (AP: −1.00 mm; ML: ±1.35 mm; DV: −4.13 mm) in Ecel1-Cre mice. The optical fibers were unilaterally implanted 100 μm above viral injection coordinates of the ZIm in Pde11a-Cre mice. The optical fibers were implanted at a 10° angle relative to the vertical plane above PAG bilaterally in Ecel1-Cre mice (AP: −4.40 mm; ML: ±1.00 mm; DV: −2.15 mm) to optogenetically stimulate axon terminals from ZIr to PAG. The optical fibers were unilaterally implanted into the PRF in Pde11a-Cre mice (AP: −4.4 mm; ML: ±0.85 mm; DV: −4.6 mm) to optogenetically stimulate axon terminals from ZIm to PRF.

To chemogenetic manipulation of GABAergic neuron subtypes, AAV2/9-hSyn-DIO-hM3Dq-mCherry or AAV2/9-hSyn-DIO-hM4Di-mCherry (80 nl) were bilaterally injected into the ZIr of Ecel1-Cre mice or ZIm of Pde11a-Cre mice. For the DTA mediated ablation experiments, AAV2/9-CAG-DIO-DTA (80nl) were bilaterally injected into the ZIr of Ecel1-Cre mice or ZIm of Pde11a-Cre mice.

### Dual color fiber photometry

Calcium signals were recorded using a dual-color fiber photometry system (Thinkertech, Nanjing Bioscience Inc.). GCamP6s was excitated by 488 channels to record changes of calcium activity, and mCherry fluorescent signal was excited by 570 nm channel as control to exclude disturbances of fluorescent signals caused by artificial factors. The laser power at the tip of the optical fiber was adjusted to be 30 μW for 488 nm excitation channel and 10 μW for the control channel (570 nm) to decrease laser bleaching of fluorescent signals. Averaged traces of Ca^2+^ fluorescent signal changes and heatmap were plotted using the MATLAB program provided by ThinkerTech Nanjing Bioscience. Adult male mice were used for dual color fiber photometry.

### *In vivo* optogenetic manipulation

Mice were recovered for at least 1 week after optical fiber implantation surgery. For optogenetic manipulation during the behavioral assessments, pulses of 465 nm laser stimulation were set by laser stimulator (INPER-B1-465, Inper, Hangzhou, China). Before behavioral tests, mice were habituated to the environments for at least 30 min after connection to a laser stimulator. The laser power at the tip of the optical fiber was adjusted to 5-10 mW for the optogenetic stimulation (10 ms pulses at 20 Hz or 5Hz). Adult male mice were used for optogenetic manipulation.

### Electrophysiology

Mice were anesthetized with pentobarbital sodium solution (50 mg/kg) via intraperitoneal injection (i.p.) and then the brains were removed and placed into ice-cold oxygenated artificial cerebrospinal fluid (ACSF) containing (in mM) 110 Choline chloride, 2.5 KCl, 0.5 CaCl_2_, 7 MgCl_2_, 1.3 NaH_2_PO_4_, 25 NaHCO_3_, 20 Glucose, 1.3 Na-Ascorbate, 0.6 Na-Pyruvate. Coronal brain slice sections (300 μm thickness) including the periaqueductal gray (PAG) or sagittal sections (250 μm thickness) including the pontine reticular formation (PRF) region were made on a vibratome (VT1200S, Leica) in ice-cold ACSF. The brain slices were incubated at 30 ± 2 ℃ for 0.5 hour and recovered for 1 hour at room temperature in a recording solution containing (in mM) 125 NaCl, 2.5 KCl, 2 CaCl_2_, 7 MgCl_2_, 1.3 NaH_2_PO_4_, 25 NaHCO_3_, 10 Glucose, 1.3 Na-Ascorbate, 0.6 Na-Pyruvate. After recovering, individual brain slices were transferred to the recording chamber and continuously perfused (2 – 4 ml/min) with ACSF containing (in mM) 125 NaCl, 2.5 KCl, 2 CaCl_2_, 7 MgCl_2_, 1.3 NaH_2_PO_4_, 25 NaHCO_3_, 10 Glucose. The incubating and recording ACSF solutions were continuously bubbled with 95% O_2_/5% CO_2_.

To test the monosynaptic and inhibitory connection of the ZIr to PAG or the ZIm to PRF, light-evoked inhibitory postsynaptic currents (eIPSCs) were recorded using pipettes (3 - 5 MΩ) filled with inner solution containing 133 CsMeSO_4_, 7 CsCl, 10 HEPES, 4 MgCl_2_, 0.5 EGTA, 3 CsOH, 4 Na_2_ATP, 0.4 Na_4_GTP, 5 QX-314 (PH 7.35; Osmolarity 310 ± 10 mOsm). And TTX (1 μM, Absin, China) was used to block the sodium channel and followed by the 4-AP (100 μM) to confirm the monosynaptic postsynaptic currents following light activation. Bicuculine (20 μM, Abcam, UK) was bath applied to validate whether the recorded IPSCs were mediated by GABA receptors. Whole-cell patch-clamp recordings were conducted with Multiclamp 700B amplifier (Axon) controlled by DigiData 1550B digitizer and analyzed with pClamp (Clampex, Clampfit, v10.7). Both male and female mice were used for electrophysiology.

### Nuclei isolation and single-nuclei RNA sequencing analysis

Sagittal brain slice sections (600 μm) from Gad67-GFP knock-in mice at postnatal day 40 (P40) were made with a vibratome as described above. The zona incerta region indicated by expression of fluorescent reporters were microdissected under a fluorescence stereomicroscope (Leica VT1200S, Germany) and collected in ice-cold artificial cerebrospinal fluid (ACSF). Two independent cohort of micro-dissected zona incerta brain slice sections (each cohort were collected from 3 male mice and 3 female mice) were harvested for single-nuclei RNA sequencing.

Nuclei isolation from micro-dissected tissue was performed using the following procedure: the collected tissue was homogenized in ice-cold RNAase-free homogenization buffer^89^ containing: 0.32 M sucrose, 5 mM CaCl_2_, 3 mM MgAC_2_, 0.1 mM EDTA (Invitrogen, AM9260G), 10 mM Tris-HCl pH 7.6 (Invitrogen, 15568-025), 0.4 U/ul Recombinant RNA inhibitor (Takara, 2312A), 0.1 mM PMSF (Roche, 10837091001), 0.1 mM β-mercaptoethanol, 1% BSA, 0.01% 10% NP40 (Thermo, 85124) in UltraPure distilled-water. Subsequently 1.5 mL 0.32 M Sucrose solution containing 0.32 M sucrose, 5 mM CaCl_2_, 3 mM MgAC_2_, 0.1 mM EDTA, 10 mM Tris-HCl pH 7.6, 0.4 U/ul Recombinant RNA inhibitor, 0.1 mM β-mercaptoethanol, 1% BSA in UltraPure distilled-water were added to the homogenates, mixed well, and centrifuged by 900 g for 10 min. After removing the supernatant, another 1.5 mL 0.32 M sucrose solution was added to the tube, resuspended, filtered by 35 μm cell-strainer (FALCON, 352235), and then centrifuged at 900 g for 8 min. The precipitate was collected and resuspended in 1.5 mL 0.32 M sucrose solution and was centrifuged at 600 g for two times. The precipitate was resuspended in 0.32 M sucrose solution and diluted with an equal volume of 50% OptiPrep density gradient medium (60% OptiPrep density gradient medium, 0.32 M sucrose solution) to give a final concentration of 25% medium solution. Slowly adding the 29% OptiPrep density gradient medium (60% OptiPrep density gradient medium, 0.32 M sucrose solution) under the 25% medium solution without disrupting the layers in a 1.5 mL centrifuge tube and centrifuged by 3000 g for 20 min^90^. After the removal of the supernatant, the nuclei pellet was dissolved in 100 μl PBS (Gibco, 10010-031). Nuclei density was analyzed using a cell counting plate (WATSON, 177-112C). Single nuclei capture was performed on 10X Genomics Single-Cell 3’ system with target capture of 8000 nuclei per sample. Single-nuclei libraries from individual samples were pulled and sequenced on the Illumina HiSeq X Ten machine. The 10x nuclei capture and library preparation protocol was performed according to the manufacturer’s recommended protocol (10X Genomics).

Sequencing reads were aligned to mouse reference genome mm10 with CellRanger (version 6.0.2). Low abundant genes (expressed in less than 3 cells) and cells of potentially low quality (nFeature (number of detected genes) < 1000 or nFeature > 6,000, or counts > 25,000, or percentage of mitochondrial genes > 5%) were removed from downstream analysis. The generated gene expression matrix files arising from two batches of experiments were further analyzed using Seurat v4 package. Briefly, SCTransform normalization was performed and RPCA was used to integrate data. 3,000 variable genes were identified by FindVariableGenes and selected to function RunPCA for the principal component analysis (PCA). Next, Uniform Manifold Approximation and Projection (UMAP) was performed to reduce variation to two dimension and FindClusters was used for unsupervised clustering of cells by setting resolution to 0.6. Subsequently, canonical cell type-specific markers were used to annotate major cell types: neurons, oligodendrocytes, endothelial cells, astrocytes, oligodendrocyte progenitor cells (OPCs), pericytes, microglia, and differentiation-committed oligodendrocyte precursors (COPs). Furthermore, we reclustered excitatory neurons and inhibitory neurons into ten and twelve subclusters, respectively. The genes differentially expressed in cells of subclusters were analyzed using FindMarkers function and used the two-sided Wilcoxon Rank-Sum test (default). Those which with *p* value < 0.05 were selected as differentially expressed genes (DEGs) candidates. Heatmap showing differentially expressed genes was generated by ggplot2 package. The regional information of neuronal subclusters were determined by the expression of DEGs in brain slices from Allen Mouse Brain Atlas (https://mouse.brain-map.org).

### Histology and immunofluorescent staining

Mice were deeply anesthetized with an intraperitoneal injection of pentobarbital sodium solution (50 mg/kg) and perfused transcardially with 0.1M phosphate-buffered saline (PBS, pH 7.4) followed by 4% paraformaldehyde (PFA). Brain tissues were collected and postfixed in 4% PFA for 6-8 hours at 4 °C. Fixed brains were then rehydrated with 28-30% sucrose in PBS at 4 °C for 48 hours. Subsequently, the brains were embedded with Optimal Cutting Temperature compound and were cut into 40 μm or 16 μm coronal sections using cryomicrotome (Thermo Scientific, CryoStar NX50). The brain slice sections were stored at −80 °C or in 0.01% sodium azide in PBS at 4 °C.

For immunofluorescent staining, brain sections were washed with 0.1M PBS for 3 times, 10 min each. Then sections were incubated with blocking buffer containing 10% donkey serum albumin (Jackson ImmunoResearch, USA) dissolved in 0.1% PBST (0.1% Triton X-100 in PBS) at room temperature for 1 hour. Sections were then incubated with primary antibody at 4 °C overnight. After washing in PBS, sections were incubated with secondary antibodies (1:1000, Invitrogen, USA) at room temperature for 1 hour. Finally, sections were mounted after counterstained with nuclear dye 4,6-diamidino-2-phenylindole (DAPI, Beyotime, China). All antibodies were titrated for working solution with blocking buffer. Primary antibody included anti-Cre recombinase (1:500, Synaptic Systems 257004, Germany) and anti-Calretinin (1:100, Swant CR6B3, Switzerland). Fluorescent images were taken with a confocal microscope (IX83-FV3000, Olympus, Japan) and a Virtual Slide Microscope VS120 (Olympus, Japan).

Fluorescence *in situ* hybridization (FISH) was performed according to the manufacture’s protocol using RNAscope multiplex fluorescent reagent kit and following designed probes were used: RNAscope® Probe-Mm-Cdh23-C2, RNAscope® Probe-Mm-Ecel1-C3, RNAscope® Probe-Mm-Ptprk-C1, RNAscope® Probe-Mm-Pde11a-C1, RNAscope® Probe-Mm-Pax6-C2, RNAscope® Probe-Mm-Nos1-C3, RNAscope® Probe-Mm-Slc32a1-C4, RNAscope® Probe-Mm-Lhx6-C2, RNAscope® Probe-Mm-SST-C4 (Advanced Cell Diagnostics, USA). The density of marker genes was calculated as the cell numbers per μm2. The distribution of marker genes was shown as the percentage of positive neuron numbers in different brain areas. Both male and female mice were used for immunofluorescent staining.

### Behavioral analysis

Self-grooming measurement. Mouse self-grooming has a complex sequenced structure that consists of several syntactic chains which beginning from elliptical bilateral paw stroke (paw and nose grooming) to unilateral strokes from the mystacial vibrissae to face following by bilateral head grooming, and finally to body licking. A 350 mm × 350 mm square box was used for the self-grooming test. Mice were placed in the chamber to acclimate for at least 10 min before the behavioral tests. For the optogenetic manipulation, mice were placed in the center of the apparatus and the behavioral assay was performed with a 3-min Laser OFF, 3-min Laser ON and 3-min Laser OFF administration pattern. In the self-grooming test when optogenetic activation of Ecel1 positive neurons in the ZIr, different frequencies (1Hz, 5Hz, 10Hz and 20 Hz) were applied to ZIr. Self-grooming behavior was mort robustly induced when optogenetic stimulating Ecel1^+^ neurons in the ZIr at 5Hz, we therefore conducted the following experiments at 5Hz. For chemogenetic manipulation, mice was intraperitoneal administrated with clozapine-N-oxide (CNO, Sigma-Aldrich; 1 mg/kg in saline for hM3Dq-mediated activation and 3 mg/kg in saline for hM4Di-mediated inhibition) 30min before self-grooming test. For water spray in chemogenetic manipulation and DTA ablation experiments, water spray was started by spraying with water toward the face of mice 4 times with a sprayer and following with a 20min recording. The total time spent in self-grooming, self-grooming bouts and duration of self-grooming per bout were measured during the following 20min.

Open field test. A 450 mm × 450 mm square box was used for the open-field test. Mice were placed in the experimental room to acclimate for at least 30 min before the behavioral tests. For the optogenetic manipulation, mice were placed in the center of the apparatus and a 12-min behavioral assay was performed with a 3-min Laser OFF and 3-min Laser ON administration pattern. The open field chamber was cleaned with 75% ethanol between each trial and the moving traces were tracked by ANY-Maze software. Total distance traveled and time spent in the center zone were analyzed.

Elevated plus maze (EPM). The EPM apparatus consisted of a central square (50 mm × 50 mm), two open arms without walls (300mm length) and two closed arms with walls (300mm length, 200mm height). For the optogenetic manipulation, mice were placed in the center of the EPM facing closed arm and a 12-min behavioral assay was performed with a 3-min Laser OFF and 3-min Laser ON administration pattern. Moving traces were tracked by ANY-Maze software. Time spent in the open arm were analyzed.

Real time place preference or aversion (RTPP). A two-chamber apparatus (500 mm × 250 mm) was used for the test. The mice were allowed to move freely in the apparatus for 10 minutes on the first day. The mice with over 80% initial place preference were excluded from the experiment. On the following two days, a 465 nm laser stimulation was delivered when mice entered the laser-coupled side for a 10min real-time laser coupling test. Moving traces were tracked by ANY-Maze software. The percentage of time in the laser-coupling side and the preference index were analyzed.

Food intake analysis. For the food intake test, mice had ad libitum access to chow in the home cage both prior to and following food intake trials. Each mouse was housed individually and its intake of normal and high-fat food was measured separately over a 24-hour period. For the optogenetic manipulation, mice were connected to a 473 nm blue laser and transferred to a transparent cage for 1-hour to habituate to the testing environment. Initially, a 10-min food intake test was conducted without any laser stimulation, followed by a 10 min trial with laser stimulation. For the 10-min trial, a 473-nm blue laser light with a pulse duration of 10-ms and varying frequencies (1Hz, 5Hz, 10Hz and 20 Hz) were applied to zona incerta, after which a 30-minute period without stimulation ensued. For the 4 ×10-min trial, a 10-min session of light stimulation (10 ms, 20 Hz) was followed by 30 min interval without stimulation, repeated four times^34^.

Novel object recognition test. The task was conducted under identical environmental conditions to those of the open-field test, consist of a training phase (also known as the familiarization phase) followed by a testing phase, with a 1-hour interval. During the training phase, mice were exposed to two objects that were identical in terms of shape, color, and odor. These objects were positioned close to the corners of one wall within the testing arena. Each mouse was placed in the center of the arena and given 10 minutes to explore the objects. In the subsequent test phase, one object was a duplicate of the one used in the training phase, while the other was a new object. Each mouse was again placed in the arena and optical stimulation was applied during the test phase, which lasted 5 minutes. The arrangement of objects during the test phase was randomly alternated between the animals within each group. The durations of time spent by the mice exploring both the familiar and the novel objects were measured.

Adult male mice were used for all the behavioral analysis above.

### EEG-EMG recordings and analysis

Electroencephalogram (EEG) recordings were made from three screws on top of the left and right cortex, at AP –3.5 mm, ML 2.5 mm and AP +1.0 mm, ML ±1.5 mm, respectively. For electromyogram (EMG) recording, two EMG electrodes (P01965, CALMONT, USA) were inserted into the neck musculature and a reference screw was implanted into the skull above the right cortex. Insulated leads from the EEG and EMG electrodes were soldered to a 2 × 3 pin header, which was secured to the skull by dental cement. For EEG and EMG recordings with fiber photometry recording or optogenetic manipulation, optical fiber was placed in the same surgery for EEG and EMG implants. After surgery, mice were allowed to recover for at least 1week before experiments. Experimental and control animals were subjected to exactly the same surgical and behavioral manipulations. All the implanted mice were housed in a 12-h dark - 12-h light cycle (Lights on between 07:00 (ZT0) and 19:00 (ZT12)). For optogenetic activation of Pde11a positive neurons in ZIm, each trial consisted of a 20 Hz pulse train (10 ms per pulse) lasting for 120 s and inter-trial interval was chosen randomly between 10 or 15 min. For chemogenetic activation or inhibition of Pde11a positive neurons in ZIm, we used the virus-transfected mice with saline as control compared to mice with CNO (1 mg/kg in saline) intraperitoneal injection. The following injection protocol was used for analysis of all mice injected with AAV virus carrying cre-dependent expression of hM3Dq or hM4Di: on day 1, we performed two consecutive saline injections at 12-h intervals at ZT5 (12:00) and ZT17 (24:00) and CNO at the same times on day 2. For DTA-mediated ablation of Pde11a positive neurons in ZIm, we recorded the virus-transfected mice for 24hr. The EEG and EMG signals were recorded and amplified using PowerLab system (AD Instruments, Australia). NREM, REM and wake states were analyzed for each 5-s epoch by LabChart8 (wake: desynchronized EEG and high EMG activity; NREM sleep: synchronized EEG with high power at 0.5–4 Hz and low EMG activity; REM sleep: desynchronized EEG with high power at theta frequencies (6–9 Hz) and low EMG activity. Experimenters were blinded to all training and behavioral assessments. Adult male mice were used for the EEG-EMG recording experiments.

### Quantification and statistical analysis

Statistical analysis was performed using Graph-Pad Prism 6, Matlab 2019a. The following statistical tests were used: two-tailed *t*-test and one-way analysis of variance (ANOVA). Data are presented as mean ± SEM. The number of cells, animals or experimental replicates are indicated in the figure legends. The animals were randomly assigned into experimental and control groups. All behavioral experiments and data analyses were conducted blindly.

## Data availability

The raw sequence data reported in this paper have been deposited in the Genome Sequence Archive in National Genomics Data Center, China National Center for Bioinformation, Beijing Institute of Genomics, Chinese Academy of Sciences (GSA: CRA017157) that are publicly accessible at https://ngdc.cncb.ac.cn/gsa. Source data are provided with this paper.

## Acknowledgement

We thank Research Assistant Zhaoxiaonan Lin from the Core Facilities of Zhejiang University School of Medicine and Dr. Sanhua Fang from the Core Facilities of Zhejiang University Institute of Neuroscience for technical support. We thank Drs. Shumin Duan, Jianhong Luo, Ying Shen (Zhejiang University) for suggestions on the project. This work was supported by grants from the Ministry of Science and Technology (2019YFA0110103 to J.C.), National Natural Science Foundation of China (81870898 to J.C.; 82090030 and 82288101 to X.-M.L.), Zhejiang Provincial Natural Science Foundation (LR18H090002), STI2030-Major Projects (2021ZD0202700 to X.-M.L.; 2021ZD0203400 to Y.-Q.Y.), Key-Area Research and Development Program of Guangdong Province (2019B030335001 to X.-M.L.), Key R&D Program of Zhejiang Province (2020C03009 and 2021C03001 to X.-M.L.), Fundamental Research Funds for the Central Universities (2021FZZX001-37 to X.-M.L.), CAMS Innovation Fund for Medical Sciences (2019-I2M-5-057 to X.-M.L.), Innovative and Entrepreneur Team of Zhejiang for 2020 Biomarker-Driven Basic and Translational Research on Major Brain Diseases (2020R01001 to X.-M.L.), National Natural Science Foundation of China Major Project (T2293733 to Y.-Q.Y.), Postdoctoral Fellowship Program of CPSF (GZC20232285 to M.Z.), and the Non-profit Central Research Institute Fund of Chinese Academy of Medical Sciences (2023-PT310-01).

## Author Contributions

J.C. conceived and designed the project, M.Z. M.W. S.L. performed in vivo circuit tracing, optogenetics, behavioral experiments and data analysis. M.Z. M.W. performed immunohistochemistry staining, fluorescence in situ hybridization and confocal imaging, J.P. H.Z. Y.Z. L.S. performed single-nuclei RNA sequencing and data analysis. X.D. performed electrophysiology experiments and data analysis. M.Z. M.W. H.Z. Y.Y. performed EEG/EMG recordings and data analysis. J.C. X.L. analyzed data and supervised the project. J.C. prepared the manuscript with inputs from all authors.

## References

1. Zeng, H. & Sanes, J.R. Neuronal cell-type classification: challenges, opportunities and the path forward. Nat Rev Neurosci 18, 530–546 (2017).

2. Zeng, H. What is a cell type and how to define it? Cell 185, 2739–2755 (2022).

3. Yao, Z., et al. A taxonomy of transcriptomic cell types across the isocortex and hippocampal formation. Cell 184, 3222–3241 e3226 (2021).

4. Tasic, B., et al. Shared and distinct transcriptomic cell types across neocortical areas. Nature 563, 72–78 (2018).

5. Yao, Z., et al. A high-resolution transcriptomic and spatial atlas of cell types in the whole mouse brain. Nature 624, 317–332 (2023).

6. DeFelipe, J., et al. New insights into the classification and nomenclature of cortical GABAergic interneurons. Nat Rev Neurosci 14, 202–216 (2013).

7. Gouwens, N.W., et al. Classification of electrophysiological and morphological neuron types in the mouse visual cortex. Nature neuroscience 22, 1182–1195 (2019).

8. Gao, L., et al. Single-neuron analysis of dendrites and axons reveals the network organization in mouse prefrontal cortex. Nat Neurosci 26, 1111–1126 (2023).

9. Cadwell, C.R., et al. Electrophysiological, transcriptomic and morphologic profiling of single neurons using Patch-seq. Nat Biotechnol 34, 199–203 (2016).

10. Fuzik, J., et al. Integration of electrophysiological recordings with single-cell RNA-seq data identifies neuronal subtypes. Nat Biotechnol 34, 175–183 (2016).

11. Gouwens, N.W., et al. Integrated Morphoelectric and Transcriptomic Classification of Cortical GABAergic Cells. Cell 183, 935–953 e919 (2020).

12. Scala, F., et al. Phenotypic variation of transcriptomic cell types in mouse motor cortex. Nature 598, 144–150 (2021).

13. Kalmbach, B.E., et al. Signature morpho-electric, transcriptomic, and dendritic properties of human layer 5 neocortical pyramidal neurons. Neuron 109, 2914–2927 e2915 (2021).

14. Zhang, M., et al. Spatially resolved cell atlas of the mouse primary motor cortex by MERFISH. Nature 598, 137–143 (2021).

15. Chen, A., et al. Spatiotemporal transcriptomic atlas of mouse organogenesis using DNA nanoball-patterned arrays. Cell 185, 1777–1792 e1721 (2022).

16. Zhang, M., et al. Molecularly defined and spatially resolved cell atlas of the whole mouse brain. Nature 624, 343–354 (2023).

17. Yao, S., et al. A whole-brain monosynaptic input connectome to neuron classes in mouse visual cortex. Nat Neurosci 26, 350–364 (2023).

18. Zhang, Z., et al. Epigenomic diversity of cortical projection neurons in the mouse brain. Nature 598, 167–173 (2021).

19. Huang, L., et al. BRICseq Bridges Brain-wide Interregional Connectivity to Neural Activity and Gene Expression in Single Animals. Cell 183, 2040 (2020).

20. Sun, Y.C., et al. Integrating barcoded neuroanatomy with spatial transcriptional profiling enables identification of gene correlates of projections. Nat Neurosci 24, 873–885 (2021).

21. Caputi, A., Melzer, S., Michael, M. & Monyer, H. The long and short of GABAergic neurons. Curr Opin Neurobiol 23, 179–186 (2013).

22. Melzer, S. & Monyer, H. Diversity and function of corticopetal and corticofugal GABAergic projection neurons. Nat Rev Neurosci 21, 499–515 (2020).

23. Malik, R., Li, Y., Schamiloglu, S. & Sohal, V.S. Top-down control of hippocampal signal-to-noise by prefrontal long-range inhibition. Cell 185, 1602–1617 e1617 (2022).

24. Lee, A.T., Vogt, D., Rubenstein, J.L. & Sohal, V.S. A class of GABAergic neurons in the prefrontal cortex sends long-range projections to the nucleus accumbens and elicits acute avoidance behavior. J Neurosci 34, 11519–11525 (2014).

25. Melzer, S., et al. Long-range-projecting GABAergic neurons modulate inhibition in hippocampus and entorhinal cortex. Science 335, 1506–1510 (2012).

26. Szabo, G.G., et al. Ripple-selective GABAergic projection cells in the hippocampus. Neuron 110, 1959–1977 e1959 (2022).

27. Schroeder, A., et al. Inhibitory top-down projections from zona incerta mediate neocortical memory. Neuron 111, 727–738 e728 (2023).

28. Saunders, A., et al. A direct GABAergic output from the basal ganglia to frontal cortex. Nature 521, 85–89 (2015).

29. Schlesiger, M.I., et al. Two septal-entorhinal GABAergic projections differentially control coding properties of spatially tuned neurons in the medial entorhinal cortex. Cell Rep 34, 108801 (2021).

30. Mitrofanis, J. Some certainty for the “zone of uncertainty“? Exploring the function of the zona incerta. Neuroscience 130, 1–15 (2005).

31. Wang, X., Chou, X.L., Zhang, L.I. & Tao, H.W. Zona Incerta: An Integrative Node for Global Behavioral Modulation. Trends Neurosci 43, 82–87 (2020).

32. Yang, Y., et al. Whole-Brain Connectome of GABAergic Neurons in the Mouse Zona Incerta. Neurosci Bull 38, 1315–1329 (2022).

33. Wang, Q., et al. The Allen Mouse Brain Common Coordinate Framework: A 3D Reference Atlas. Cell 181, 936–953 e920 (2020).

34. Zhang, X. & van den Pol, A.N. Rapid binge-like eating and body weight gain driven by zona incerta GABA neuron activation. Science 356, 853–859 (2017).

35. Liu, K., et al. Lhx6-positive GABA-releasing neurons of the zona incerta promote sleep. Nature 548, 582–587 (2017).

36. Shang, C., et al. A subcortical excitatory circuit for sensory-triggered predatory hunting in mice. Nat Neurosci 22, 909–920 (2019).

37. Zhao, Z.D., et al. Zona incerta GABAergic neurons integrate prey-related sensory signals and induce an appetitive drive to promote hunting. Nat Neurosci 22, 921–932 (2019).

38. Ogasawara, T., et al. A primate temporal cortex-zona incerta pathway for novelty seeking. Nat Neurosci 25, 50–60 (2022).

39. Venkataraman, A., et al. Modulation of fear generalization by the zona incerta. Proc Natl Acad Sci U S A 116, 9072–9077 (2019).

40. Li, Z., Rizzi, G. & Tan, K.R. Zona incerta subpopulations differentially encode and modulate anxiety. Sci Adv 7, eabf6709 (2021).

41. Ahmadlou, M., et al. A cell type-specific cortico-subcortical brain circuit for investigatory and novelty-seeking behavior. Science 372 (2021).

42. Lin, S., et al. Somatostatin-Positive Neurons in the Rostral Zona Incerta Modulate Innate Fear-Induced Defensive Response in Mice. Neurosci Bull (2022).

43. Velmeshev, D., et al. Single-cell genomics identifies cell type-specific molecular changes in autism. Science 364, 685–689 (2019).

44. Huang, K.W., et al. Molecular and anatomical organization of the dorsal raphe nucleus. Elife 8 (2019).

45. Zeisel, A., et al. Molecular Architecture of the Mouse Nervous System. Cell 174, 999–1014 e1022 (2018).

46. La Manno, G., et al. Molecular Diversity of Midbrain Development in Mouse, Human, and Stem Cells. Cell 167, 566–580 e519 (2016).

47. Kee, N., et al. Single-Cell Analysis Reveals a Close Relationship between Differentiating Dopamine and Subthalamic Nucleus Neuronal Lineages. Cell Stem Cell 20, 29–40 (2017).

48. Chen, R., Wu, X., Jiang, L. & Zhang, Y. Single-Cell RNA-Seq Reveals Hypothalamic Cell Diversity. Cell Rep 18, 3227–3241 (2017).

49. Zhang, Y.H., et al. Cascade diversification directs generation of neuronal diversity in the hypothalamus. Cell Stem Cell 28, 1483–1499 e1488 (2021).

50. Marques, S., et al. Oligodendrocyte heterogeneity in the mouse juvenile and adult central nervous system. Science 352, 1326–1329 (2016).

51. Lein, E.S., et al. Genome-wide atlas of gene expression in the adult mouse brain. Nature 445, 168–176 (2007).

52. Xu, Y., et al. Identification of a neurocircuit underlying regulation of feeding by stress-related emotional responses. Nat Commun 10, 3446 (2019).

53. Mu, M.D., et al. A limbic circuitry involved in emotional stress-induced grooming. Nat Commun 11, 2261 (2020).

54. Gao, Z.R., et al. Tac1-Expressing Neurons in the Periaqueductal Gray Facilitate the Itch-Scratching Cycle via Descending Regulation. Neuron 101, 45–59 e49 (2019).

55. Xie, Z., et al. A brain-to-spinal sensorimotor loop for repetitive self-grooming. Neuron 110, 874–890 e877 (2022).

56. Zhang, Y.F., et al. Ventral striatal islands of Calleja neurons control grooming in mice. Nat Neurosci 24, 1699–1710 (2021).

57. Kalueff, A.V., et al. Neurobiology of rodent self-grooming and its value for translational neuroscience. Nat Rev Neurosci 17, 45–59 (2016).

58. Sun, J., et al. Excitatory SST neurons in the medial paralemniscal nucleus control repetitive self-grooming and encode reward. Neuron 110, 3356–3373 e3358 (2022).

59. Mangieri, L.R., et al. A neural basis for antagonistic control of feeding and compulsive behaviors. Nat Commun 9, 52 (2018).

60. Liu, Q., et al. An iterative neural processing sequence orchestrates feeding. Neuron 111, 1651–1665 e1655 (2023).

61. Ahnaou, A., et al. Long-term enhancement of REM sleep by the pituitary adenylyl cyclase-activating polypeptide (PACAP) in the pontine reticular formation of the rat. Eur J Neurosci 11, 4051–4058 (1999).

62. Maloney, K.J., Mainville, L. & Jones, B.E. c-Fos expression in GABAergic, serotonergic, and other neurons of the pontomedullary reticular formation and raphe after paradoxical sleep deprivation and recovery. J Neurosci 20, 4669–4679 (2000).

63. Yamahachi, H., Marik, S.A., McManus, J.N., Denk, W. & Gilbert, C.D. Rapid axonal sprouting and pruning accompany functional reorganization in primary visual cortex. Neuron 64, 719–729 (2009).

64. Bagri, A., Cheng, H.J., Yaron, A., Pleasure, S.J. & Tessier-Lavigne, M. Stereotyped pruning of long hippocampal axon branches triggered by retraction inducers of the semaphorin family. Cell 113, 285–299 (2003).

65. Luo, L. & O’Leary, D.D. Axon retraction and degeneration in development and disease. Annu Rev Neurosci 28, 127–156 (2005).

66. Wen, S., et al. Spatiotemporal single-cell analysis of gene expression in the mouse suprachiasmatic nucleus. Nat Neurosci 23, 456–467 (2020).

67. Kim, D.W., et al. Gene regulatory networks controlling differentiation, survival, and diversification of hypothalamic Lhx6-expressing GABAergic neurons. Commun Biol 4, 95 (2021).

68. Puelles, L., et al. LacZ-reporter mapping of Dlx5/6 expression and genoarchitectural analysis of the postnatal mouse prethalamus. J Comp Neurol 529, 367–420 (2021).

69. Wang, X.Y., et al. A cross-modality enhancement of defensive flight via parvalbumin neurons in zonal incerta. Elife 8 (2019).

70. Kalueff, A.V. & Tuohimaa, P. Grooming analysis algorithm for neurobehavioural stress research. Brain Res Brain Res Protoc 13, 151–158 (2004).

71. Kulkosky, P.J., Gibbs, J. & Smith, G.P. Feeding suppression and grooming repeatedly elicited by intraventricular bombesin. Brain Res 242, 194–196 (1982).

72. Krahn, D.D., Gosnell, B.A., Grace, M. & Levine, A.S. CRF antagonist partially reverses CRF- and stress-induced effects on feeding. Brain Res Bull 17, 285–289 (1986).

73. Rodrigo-Angulo, M.L., Heredero, S., Rodriguez-Veiga, E. & Reinoso-Suarez, F. GABAergic and non-GABAergic thalamic, hypothalamic and basal forebrain projections to the ventral oral pontine reticular nucleus: their implication in REM sleep modulation. Brain Res 1210, 116–125 (2008).

74. Vanini, G., Wathen, B.L., Lydic, R. & Baghdoyan, H.A. Endogenous GABA levels in the pontine reticular formation are greater during wakefulness than during rapid eye movement sleep. J Neurosci 31, 2649–2656 (2011).

75. Xi, M.C., Morales, F.R. & Chase, M.H. Evidence that wakefulness and REM sleep are controlled by a GABAergic pontine mechanism. J Neurophysiol 82, 2015–2019 (1999).

76. Camacho-Arroyo, I., Alvarado, R., Manjarrez, J. & Tapia, R. Microinjections of muscimol and bicuculline into the pontine reticular formation modify the sleep-waking cycle in the rat. Neurosci Lett 129, 95–97 (1991).

77. Blanco-Centurion, C., Vidal-Ortiz, A., Sato, T. & Shiromani, P.J. Activity of GABA neurons in the zona incerta and ventral lateral periaqueductal grey is biased towards sleep. Sleep 46 (2023).

78. Chen, K.S., et al. A Hypothalamic Switch for REM and Non-REM Sleep. Neuron 97, 1168–1176 e1164 (2018).

79. Weber, F., et al. Control of REM sleep by ventral medulla GABAergic neurons. Nature 526, 435–438 (2015).

80. Xu, M., et al. Basal forebrain circuit for sleep-wake control. Nat Neurosci 18, 1641–1647 (2015).

81. Pinto, L., et al. Fast modulation of visual perception by basal forebrain cholinergic neurons. Nat Neurosci 16, 1857–1863 (2013).

82. Liu, D. & Dan, Y. A Motor Theory of Sleep-Wake Control: Arousal-Action Circuit. Annu Rev Neurosci 42, 27–46 (2019).

83. McGinley, M.J., et al. Waking State: Rapid Variations Modulate Neural and Behavioral Responses. Neuron 87, 1143–1161 (2015).

84. John, J., Wu, M.F., Boehmer, L.N. & Siegel, J.M. Cataplexy-active neurons in the hypothalamus: implications for the role of histamine in sleep and waking behavior. Neuron 42, 619–634 (2004).

85. Plaha, P., Ben-Shlomo, Y., Patel, N.K. & Gill, S.S. Stimulation of the caudal zona incerta is superior to stimulation of the subthalamic nucleus in improving contralateral parkinsonism. Brain 129, 1732–1747 (2006).

86. Mostofi, A., et al. Outcomes from deep brain stimulation targeting subthalamic nucleus and caudal zona incerta for Parkinson’s disease. NPJ Parkinsons Dis 5, 17 (2019).

87. Summa, K.C., et al. Disrupted sleep-wake regulation in the MCI-Park mouse model of Parkinson’s disease. NPJ Parkinsons Dis 10, 54 (2024).

88. Tandberg, E., Larsen, J.P. & Karlsen, K. A community-based study of sleep disorders in patients with Parkinson’s disease. Mov Disord 13, 895–899 (1998).

89. Corces, M.R., et al. An improved ATAC-seq protocol reduces background and enables interrogation of frozen tissues. Nat Methods 14, 959–962 (2017).

90. Tuesta, L.M., et al. In vivo nuclear capture and molecular profiling identifies Gmeb1 as a transcriptional regulator essential for dopamine neuron function. Nat Commun 10, 2508 (2019).

