## Supplemental Material for "Transcriptomic and spatial GABAergic neuron subtypes in zona incerta mediate distinct innate behaviors"

### Supplementary Figures and Figure Legends

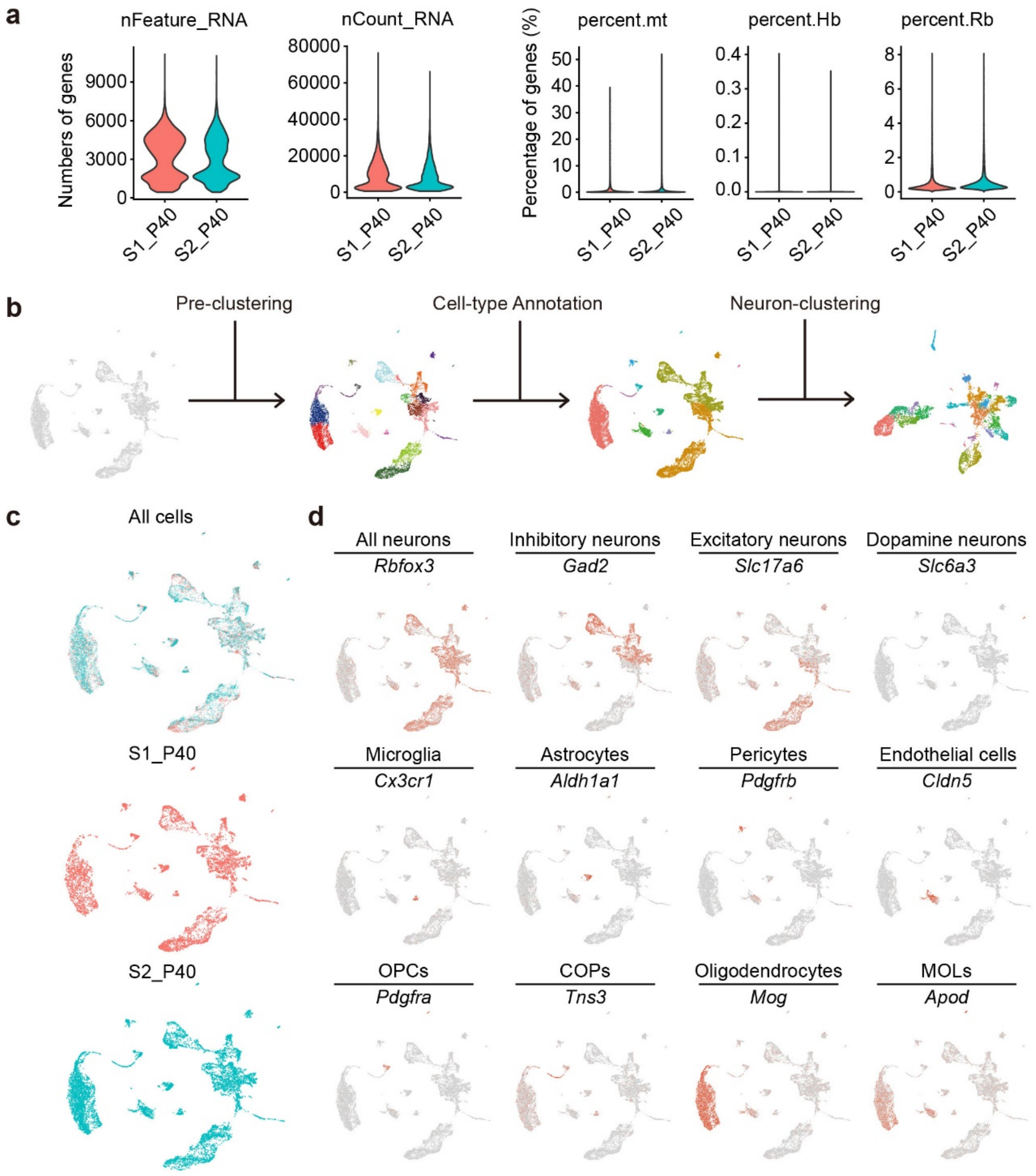


**Supplementary Figure 1: Single- nucleus RNA sequencing dataset quality matrix and data analysis.**

**a,** Violin plots showing the number of genes (nFeature), unique molecular identifiers (nCount), percent of mitochondria genes (percent.mt), percent of hemoglobin genes (percent.Hb) and ribosome genes (percent.Rb) detected in two cohorts (S1_P40, S2_P40) of 10x genomics single-nucleus RNA-sequencing (snRNA-seq) experiment before custom filtering, respectively.

**b,** Workflow of clustering analysis of total snRNA-seq cells or neurons.

**c,** UMAP plot showing all cell clusters of total snRNA-seq cells and each snRNA-seq replicate of the two independent cohorts (S1_P40 in orange and S2_P40 in green).

**d,** Feature plots depicting example marker genes used to annotate the major cell types. OPCs, oligodendrocyte precursor cells; COPs, committed oligodendrocyte precursors; MOLs, mature oligodendrocytes. Source data are provided as a Source Data file.

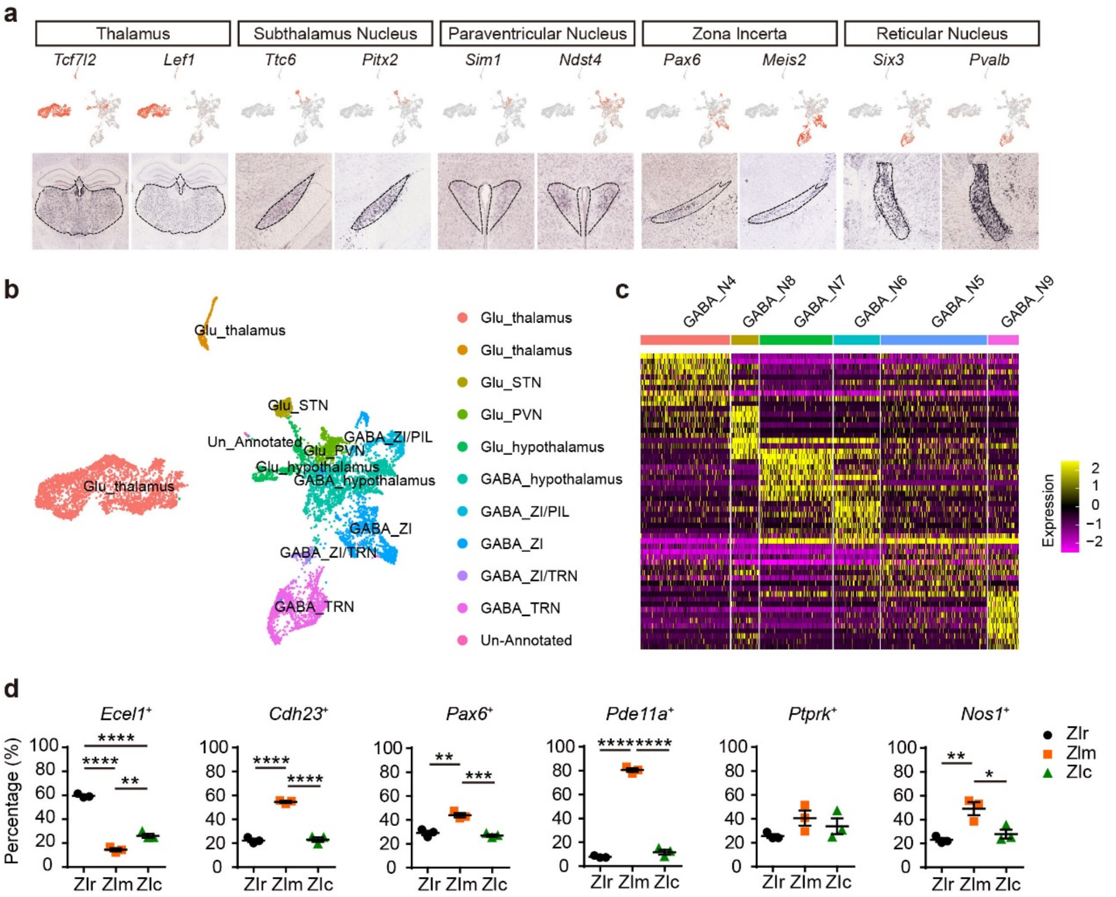


**Supplementary Figure 2: Annotation of regional distribution of glutamatergic and GABAergic neuron clusters.**

**a,** *In situ* hybridization data (from Allen Brain Atlas, https://mouse.brain-map.org/) showing the expression of region specific DEGs in mouse brain slice sections. Note that coronal section images were used for all genes except for Six3 and Pvalb in the reticular nucleus (sagittal sections).

**b,** Annotation of regional distribution of glutamatergic and GABAergic neuron clusters.

**c,** Heatmap showing the DEGs of six GABAergic neuron clusters in zona incerta.

**d,** Statistical results showing the expression and distribution of *Ecel1* (ZIr, 59.56 ± 0.93%; ZIm, 14.41 ± 1.33%; ZIc, 26.03 ± 1.76%; ****$P < 0.0001$, **$P = 0.0025$), *Cdh23* (ZIr, 22.37 ± 1.40%; ZIm, 54.60 ± 0.89%; ZIc, 23.03 ± 1.87%; ****$P <$ 0.0001), *Pax6* (ZIr, 29.20 ± 1.83%; ZIm, 43.97 ± 1.79%; ZIc, 26.83 ± 1.02%; **$P = 0.0015$, ***$P = 0.0007$), *Pde11a* (ZIr, 7.873 ± 0.55%; ZIm, 80.49 ± 1.57%; ZIc, 11.63 ± 2.04%; ****$P < 0.0001$), *Ptprk* (ZIr, 25.59 ± 1.67%; ZIm, 40.59 ± 6.35%; ZIc, 33.82 ± 6.63%; $P = 0.2011$, ZIr vs ZIm; $P = 0.5599$, ZIr vs ZIc; $P =$ 0.6665, ZIm vs ZIc), *Nos1* (ZIr, 22.98 ± 1.68%; ZIm, 49.20 ± 5.38%; ZIc, 27.83 ± 3.71%; **$P = 0.0075$, *$P = 0.0192$) in the ZIr, ZIm and ZIc, n = 9 brain slices from 3 mice per group. One-way ANOVA and tukey's multiple comparisons test. Data are presented as mean ± SEM. Source data are provided as a Source Data file.

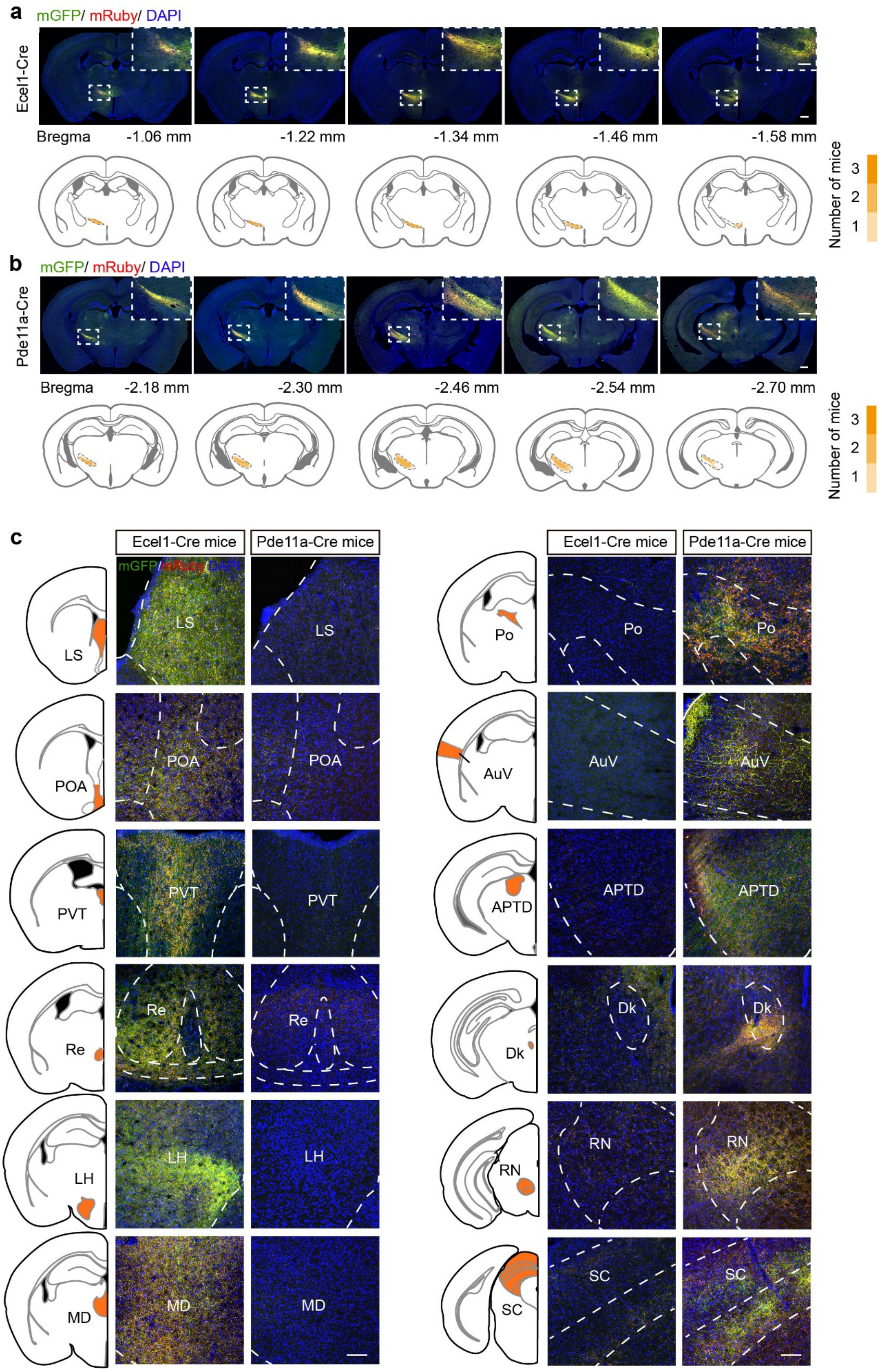


**Supplementary Figure 3: Different target brain regions of efferent projections from ZIr$^{Ecel1}$ and ZIm$^{Pde11a}$ neurons.**

**a,** Representative images showing the overlay of mGFP and synaptophysin-mRuby expression from anterior to posterior brain slice sections following injection of AAV virus in the ZIr of Ecel1-cre mice (**a**) or the ZIm of Pde11a-cre mice (**b**). Scale bar, 500 μm and 250 μm (zoom-in image). Yellow shaded areas indicate the range of viral infection at the injection site.

**c,** Representative images of axon terminals from $\mathrm{ZIr}^{\mathrm{Ecel1}}$ neurons and $\mathrm{ZIm}^{\mathrm{Pde11a}}$ neurons in downstream areas. Scale bar, 100 μm. Yellow shaded areas indicate the expression area of neuronal terminals. The experiment was independently repeated 3 times with similar results for each mouse strain. Abbreviations: LS, lateral septal nucleus; POA, preoptic area; PVT, paraventricular thalamic nucleus; Re, reuniens thalamic nucleus; LH, lateral hypothalamic area; MD, mediodorsal thalamic nucleus; Po, posterior thalamic nucleus; Auv, secondary auditory cortex; ventral area; APTD, anterior pretectal nucleus; Dk, nucleus of Darkschewitsch; RN, red nucleus; SC, superior colliculus.

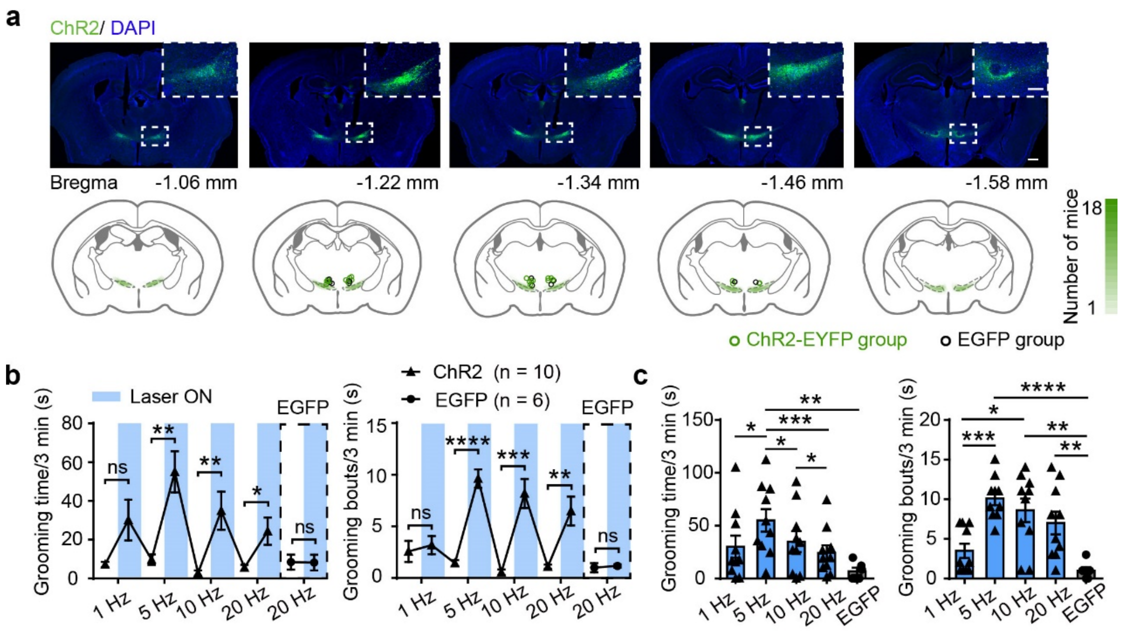


**Supplementary Figure 4: Optogenetic activation of $\mathrm{ZIr}^{\mathrm{Ecel1}}$ neurons in different frequencies.**

**a,** Representative images showing the overlay of ChR2-EYFP or EGFP expression from anterior to posterior brain slice sections from Ecel1-cre mice. Lower panel: the location of fiber placement for optogenetic stimulation in ZIr

(Green circles: ChR2-EYFP group, n = 12. Black circles: EGFP group, n = 6). Scale bar, 500 μm and 250 μm (zoom-in image). Green shaded areas indicate the range of viral infection at the injection site.

**b,** The time spent on self-grooming (left) and grooming bouts (right) in 3min at different frequencies of optogenetic stimulation in Ecel1-Cre mice. Grooming time: 5Hz, ***P* = 0.0019, 10Hz, ***P* = 0.0069, 20Hz, **P* = 0.0265. Grooming bouts: 5Hz, *****P* < 0.0001, 10Hz, ****P* = 0.0004, 20Hz, ***P* = 0.0018. Dashed frame indicates the EGFP-group. Blue column indicates laser stimulation period (3 min).

**c,** The time spent on self-grooming (left) and grooming bouts (right) in 3min at different frequencies during optogenetic stimulation of $ZIr^{Ecel1}$ neurons. Grooming time: 1Hz *vs* 5Hz, **P* = 0.0317; 5Hz *vs* 10Hz, **P* = 0.0250; 5Hz *vs* 20Hz, ****P* = 0.0008; 10Hz *vs* 20Hz, **P* = 0.0452; 5Hz *vs* EGFP, ***P* = 0.0042. Grooming bouts: 1Hz *vs* 5Hz, ****P* = 0.0007; 1Hz *vs* 10Hz, **P* = 0.0168; 5Hz *vs* EGFP, *****P* < 0.0001; 10Hz *vs* EGFP, ***P* = 0.0018; 20Hz *vs* EGFP, ***P* = 0.007. Two-tailed paired t test for (**b**) and (**c**). Two-tailed unpaired t test for (**c**). Data are presented as mean ± SEM. Source data are provided as a Source Data file.

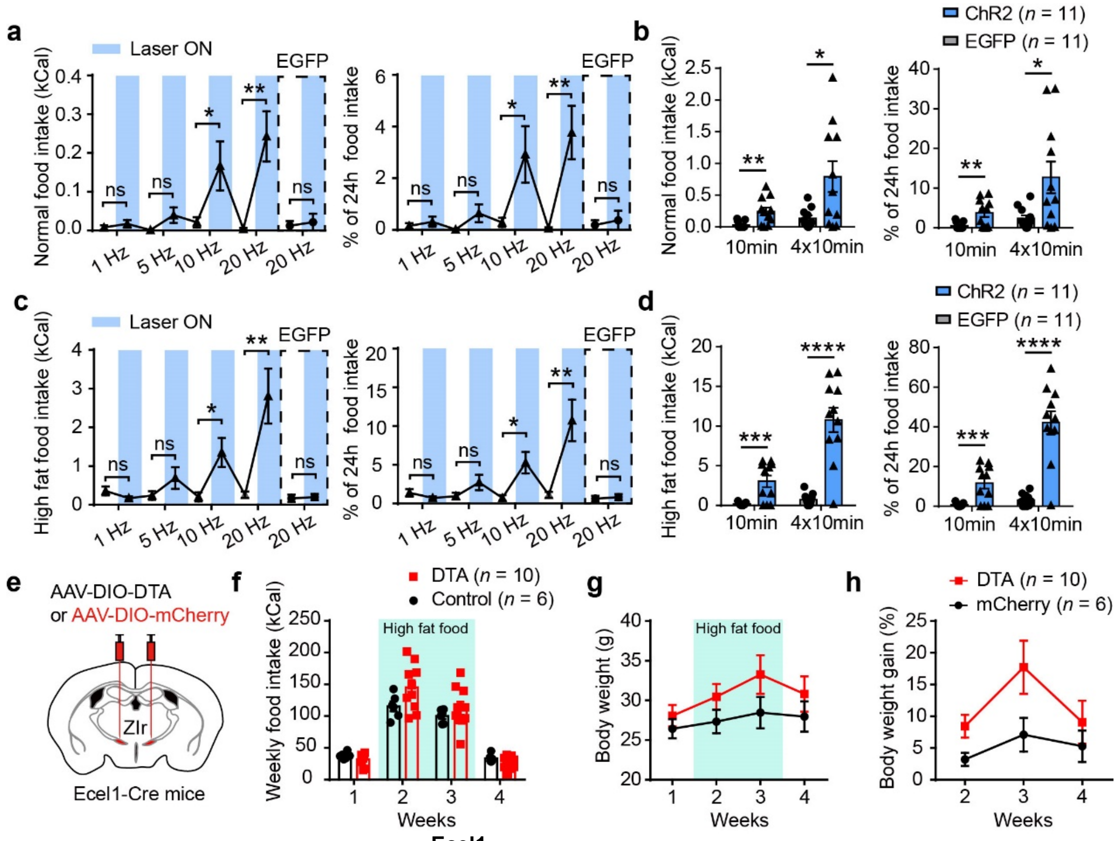


**Supplementary Figure 5: ZIr$^{Ecel1}$ neurons in food intake test.**

**a,** Normal food intake (left) and the percentage of unstimulated 24-hour intake (right) at different frequencies of optogenetic stimulation in Ecel1-Cre mice. Normal food intake: 10Hz, *$P$ = 0.0408, 20Hz, **$P$ = 0.004. Percentage of unstimulated 24-hour food intake, 10Hz, *$P$ = 0.0354, 20Hz, **$P$ = 0.005. Dashed frame indicates the EGFP-group. Blue column indicates laser stimulation period (3 min).

**b,** Normal food intake (left) and the percentage of unstimulated 24-hour food intake (right) during 10 min and four times 10 min from ChR2-EYFP group and EGFP group mice at light stimulation (10 ms, 20 Hz). Normal food intake, **$P$ = 0.0045, *$P$ = 0.0154. Percentage of unstimulated 24-hour food intake, **$P$ = 0.0061, *$P$ = 0.0196.

**c,** High-fat food intake (left) and the percentage of unstimulated 24-hour intake (right) at different frequencies of optogenetic stimulation in ChR2-EYFP group mice. High-fat food intake: 10Hz, *$P$ = 0.0285, 20Hz, **$P$ = 0.0044. Percentage of unstimulated 24-hour food intake, 10Hz, *$P$ = 0.0208, 20Hz, **$P$ = 0.0043. Dashed frame indicates the EGFP-group. Blue column indicates laser stimulation period (3 min).

**d,** High-fat food intake (left) and the percentage of unstimulated 24-hour food intake (right) during 10 min and four times 10 min from ChR2-EYFP group and EGFP group mice at light stimulation (10 ms, 20 Hz). High fat food intake, ***$P$ = 0.0004, ****$P$ < 0.0001. Percentage of unstimulated 24-hour food intake, ***$P$ = 0.0004, ****$P$ < 0.0001.

**e,** Schematic diagram showing the injection of AAV-DIO-DTA to ablate the ZIr$^{Ecel1}$ neurons of Ecel1-Cre mice.

**f,** Weekly food intake (**f**) and body weight (**g**) from DTA and mCherry group mice. The green bar indicates the high fat food given in the 2 and 3 weeks.

**h,** Percentage of body weight gain in the 2-4 weeks compared to first week of mice from the DTA and mCherry group.

Two-tailed paired t test for (**a**) and (**c**). Two-tailed unpaired t test for (**b**), (**d**). Two-way ANOVA and Sidak's multiple comparisons test for (**f-h**). Data are presented as mean ± SEM. Source data are provided as a Source Data file.

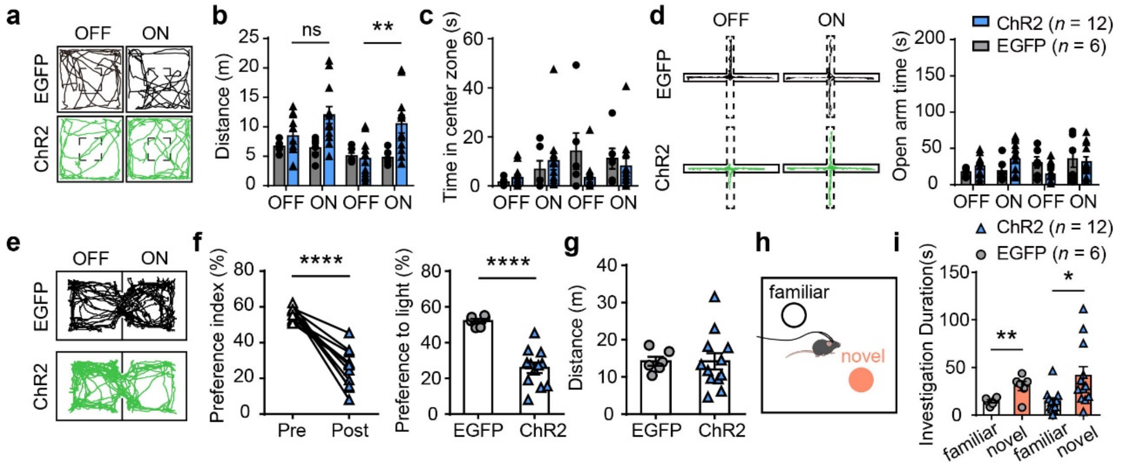


**Supplementary Figure 6: Effects of activating ZIr$^{Ecel1}$ neurons in different behavioral paradigms.**

**a,** Representative traces showing the movement of Ecel1-cre mice upon optogenetic stimulation of ZIr$^{Ecel1}$ neurons in open field test (Black: EGFP group. Green: ChR2-EYFP group).

**b,** Statistical analysis of locomotion distance (**b**) and time spent in center zone (**c**) of Ecel1-cre mice upon optogenetic stimulation of ChR2-EYFP or EGFP group, respectively. Distance: **P* = 0.0123, **P* = 0.0109 (ChR2 *vs* EGFP). *P* = 0.1389, ***P* = 0.0043 (ChR2 ON *vs* ChR2 OFF). Time spent in center zone, *P* = 0.5924.

**d,** Representative traces and statistical analysis of ChR2-EYFP or EGFP group in the elevated plus maze test of Ecel1-cre mice upon optogenetic stimulation of $ZIr^{Ecel1}$ neurons.

**e-g,** Representative traces (**e**) and statistical analysis (**f** and **g**) of ChR2-EYFP or EGFP group in the real-time place preference test of Ecel1-cre mice upon optogenetic stimulation of $ZIr^{Ecel1}$ neurons. Preference index (**f**), *****P* < 0.0001. Preference to light (**g**), *****P* < 0.0001.

**h,** Schematics of novel object recognition test with a familiar object and a novel object.

**i,** The statistical analysis show the duration of investigation of Ecel1-cre mice in 5min upon optogenetic stimulation of ChR2-EYFP or EGFP group. EGFP group, ***P* = 0.0076. ChR2-EYFP group, **P* = 0.0117.

Two-tailed paired t test for (**f**) and (**i**). Two-tailed unpaired *t*-test for (**f**) and (**g**). One way ANOVA and Tukey's multiple comparisons test for (**b**). Two-way ANOVA with Sidak's multiple comparisons test for (**b**), (**c**) and (**d**). Data are presented as mean ± SEM. Source data are provided as a Source Data file.

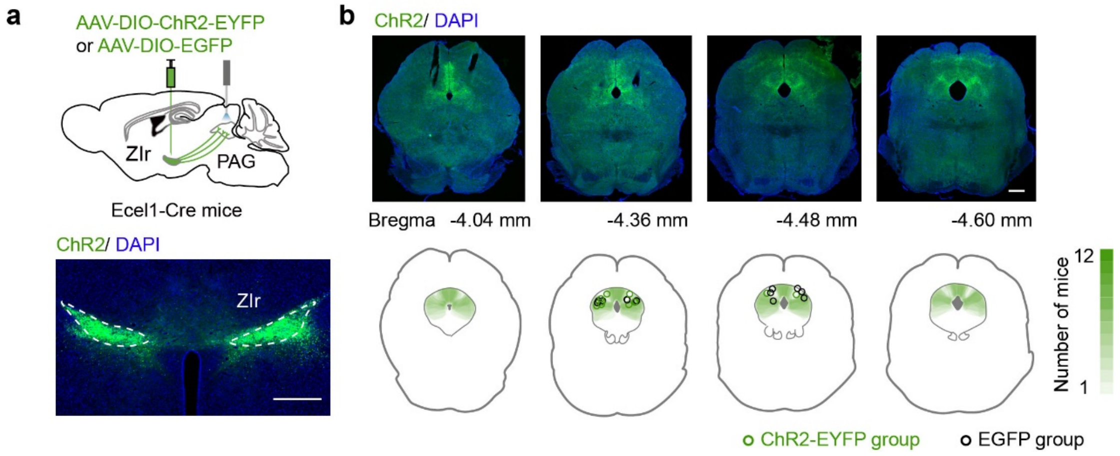


**Supplementary Figure 7: The expression of ChR2-EYFP terminals and location of optical fiber in PAG of Ecel1-Cre mice.**

**a,** Schematic diagram and representative images showing the expression of ChR2-EYFP in ZIr and optical fiber planted above the PAG of the Ecel1-cre mice. Scale bar, 500 μm. The experiment was independently repeated 6 times with similar results for each mouse strain.

**b,** The expression of ChR2-EYFP axon terminals of $ZIr^{Ecel1}$ neurons from anterior to posterior brain slices sections in PAG. Lower panel, schematic showing overlay of location of optical fiber placement in PAG with ChR2-EYFP or EGFP expression. Green circles (ChR2-EYFP group, n = 6) and black circles (EGFP group, n = 6) indicate location of fiber placement for optogenetic stimulation in PAG. Scale bar, 500 μm. Green shaded areas indicate the expression range of neuronal terminals.

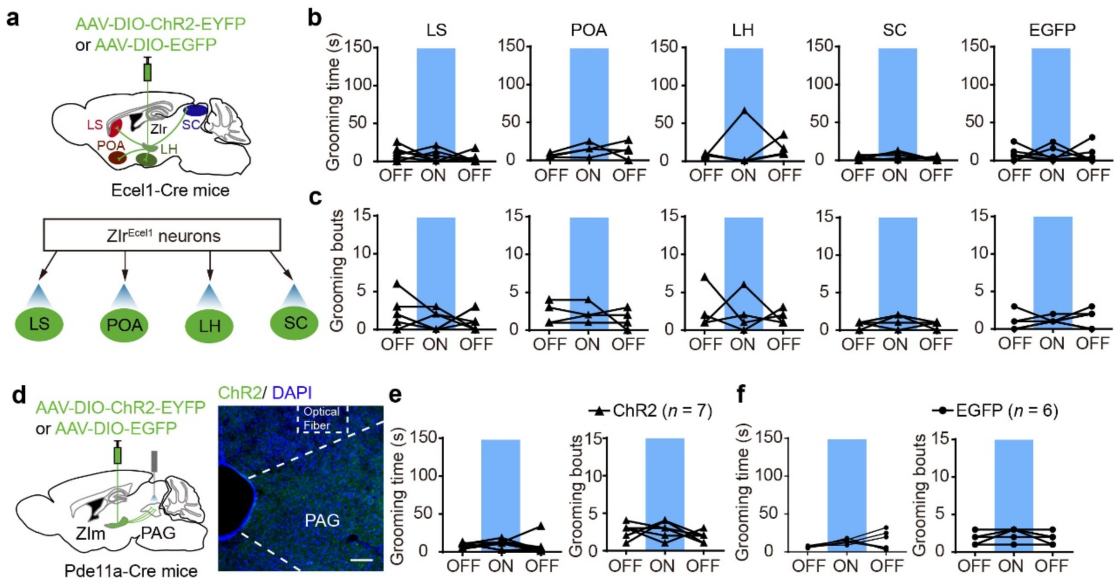


**Supplementary Figure 8: Activation of ZIr$^{Ecel1}$ neurons to other targets or ZIm$^{Pde11a}$ neurons to PAG pathway had no effects on self-grooming.**

**a,** Schematic diagram showing optogenetic stimulation of the ChR2-EYFP or EGFP expressing terminals from ZIr$^{Ecel1}$ to LS, POA, LH or SC pathway in Ecel1-Cre mice.

**b,** Statistic analysis showing time spent for self-grooming (**b**) and grooming bouts (**c**) in 3 min before, during or after optogenetic stimulation of ZIr$^{Ecel1}$ terminals in the LS, POA, LH or SC in ChR2-EYFP group and EGFP group, respectively. N = 6 mice for ZIr$^{Ecel1}$-LS group, n = 4 mice for ZIr$^{Ecel1}$-POA group, n = 4 mice for ZIr$^{Ecel1}$-LH group, n = 5 mice for ZIr$^{Ecel1}$-SC group, n = 6 mice for EGFP group.

**d,** Schematic diagram showing optogenetic stimulation of the ChR2-EYFP or EGFP expressing terminals from ZIm$^{Pde11a}$ to PAG pathway in Pde11a-Cre mice. The experiment was independently repeated 7 times with similar results for each mouse strain.

**e,** Statistic analysis showing time spent for self-grooming (left) and grooming bouts (right) in 3 min before, during or after optogenetic stimulation of ZIm$^{Pde11a}$ to PAG pathway in ChR2-EYFP group (**e**) and EGFP group (**f**).

One-way ANOVA for (**b**), (**c**), (**e**) and (**f**). Blue column indicates laser stimulation period (20 Hz, 3min).

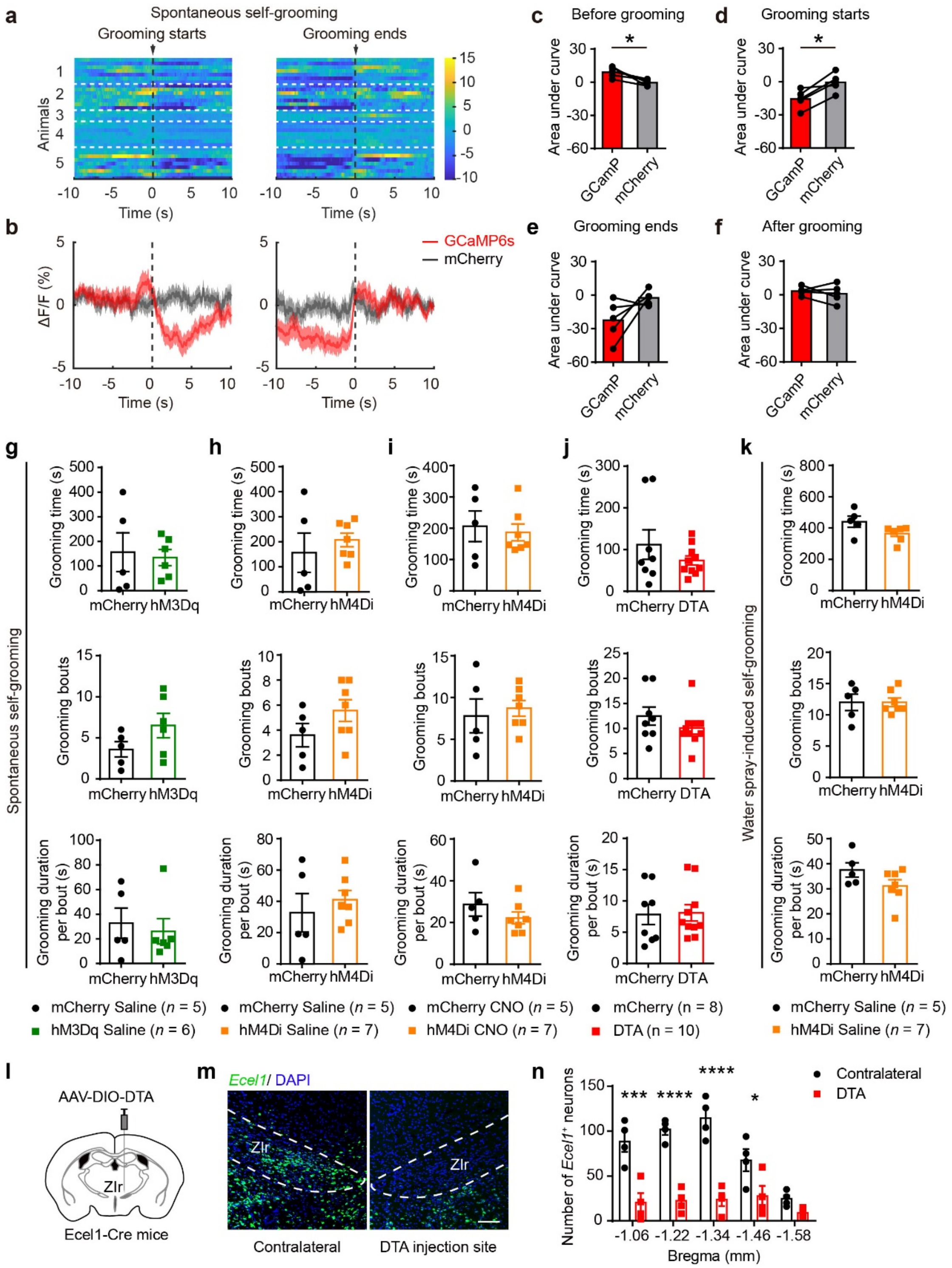


**Supplementary Figure 9: ZIr$^{Ecel1}$ neurons participate in regulating spontaneous and water-spray induced self-grooming.**

**a,** Average calcium transients (**a**) and heatmap (**b**) of ZIr$^{Ecel1}$ neurons before, during and after spontaneous self-grooming. Red trace: recording of GCaMP6s fluorescent signal; Gray trace: recording of control mCherry fluorescent signal. The dashed lines indicate the start or the end of self-grooming. Shaded areas around means indicate standard error of mean (SEM).

**c-f,** Statistic analysis showing area under curve (AUC) of average calcium transients of GCamP6s channel compared with mCherry control channel before self-grooming starts (**c**, **P* = 0.0199) and after self-grooming starts (**d**, **P* = 0.0104), before grooming ends (**e**, *P* = 0.1192) and after grooming ends (**f**, *P* = 0.5212) in spontaneous self-grooming. N = 5 mice per group.

**g-j,** Statistic analysis of time spent for spontaneous self-grooming (upper), grooming bouts (mid) and grooming duration per bout (lower) in 20 min following injection of saline in hM3Dq-group mice (**g**) and hM4Di-group mice (**h**) or injection of CNO in hM4Di-group mice (**i**) or DTA-group mice (**j**) compared with mCherry-group mice.

**k,** Statistic analysis of time spent for water spray induced self-grooming (upper), grooming bouts (mid) and grooming duration per bout (lower) in 20min following injection of saline in hM4Di-group mice compared with mCherry-group mice.

**l-n,** Representative images (**m**) and statistical results (**n**) showing *Ecel1*$^+$ neuronal loss in the ZIr from mice with AAV-DIO-DTA unilateral injection in the ZIr. ****P* = 0.0003, *****P* < 0.0001, **P* = 0.0292, n=4 mice. Scale bar, 100 μm. The experiment was independently repeated 4 times with similar results for each mouse strain.

Two-tailed paired *t* test for (**c-f**). Two-tailed unpaired *t* test for (**g-k**). Two-way ANOVA and Sidak's multiple comparisons test for (**n**). Data are presented as mean ± SEM. Source data are provided as a Source Data file.

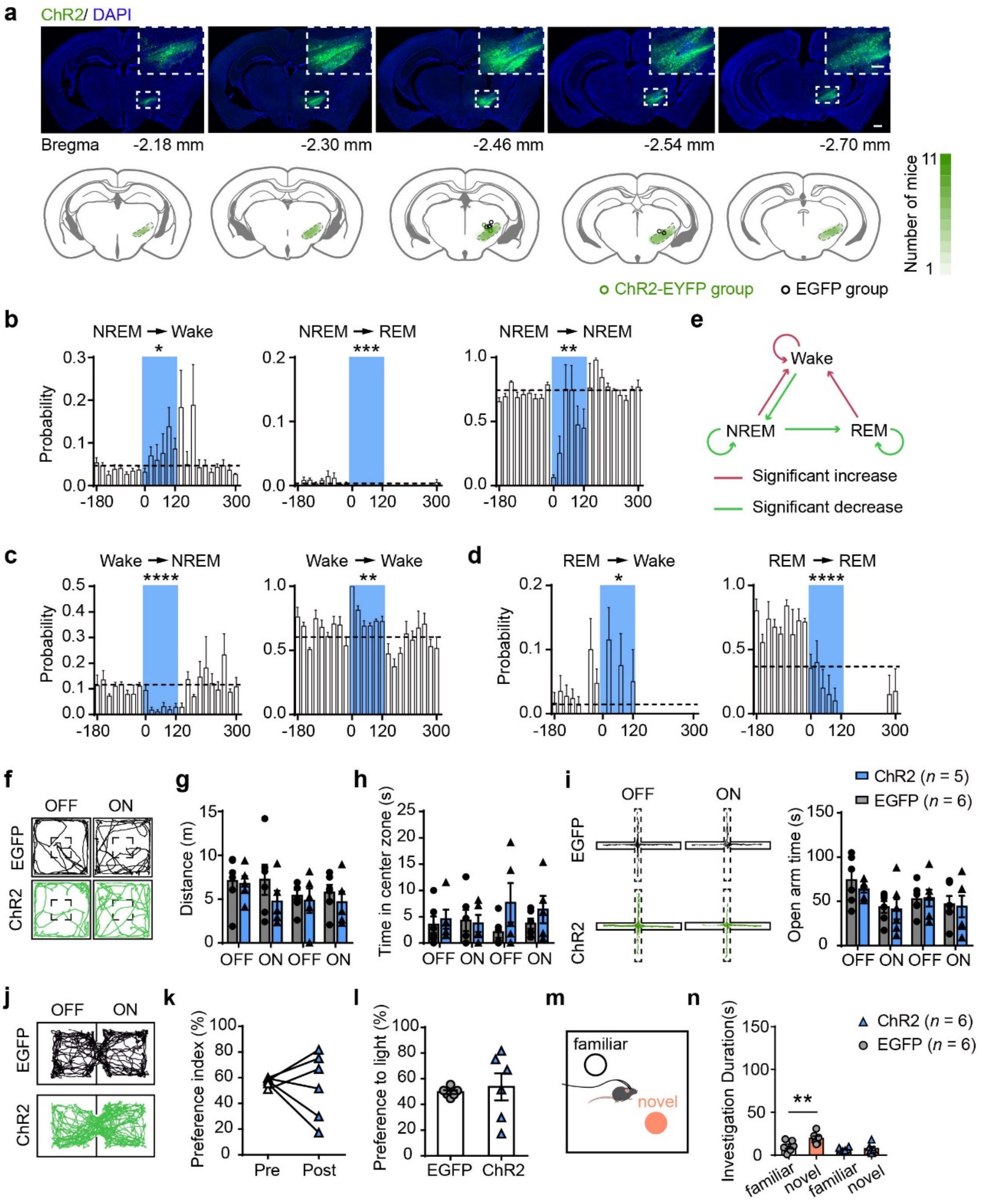


**Supplementary Figure 10: Optogenetic activation of ZIm$^{Pde11a}$ neurons significantly increases transition probabilities from sleep to wakefulness.**

**a,** Representative images showing the overlay of ChR2-EYFP or EGFP expression from anterior to posterior slice sections from Pde11a-cre mice. Lower panel: the location of fiber placement for optogenetic stimulation in ZIm (Green circles: ChR2-EYFP group, n = 5. Black circles: EGFP group, n = 6). Scale bar, 500 μm and 250 μm (zoom-in image). Green shaded areas indicate the range of viral infection at the injection site.

**b-d,** Transition probability between each pair of brain states during optogenetic light stimulation. Each bin represents transition probabilities within each 20 s period. Dashed line, baseline transition probability. Blue column indicates laser stimulation period (20 Hz, 120 s). *$P$ < 0.05, **$P$ < 0.005, ***$P$ = 0.0005 ****$P$ < 0.0001, n = 5 mice.

**e,** Schematic diagram showing transition probabilities between different brain states. Red, significant increase. Green, significant decrease.

**f,** Representative traces showing the movement of Pde11a-cre mice upon optogenetic stimulation of $ZIm^{Pde11a}$ neurons in open field test. Black trace: EGFP group. Green trace: ChR2-EYFP group.

**g-h,** Statistical analysis of locomotion distance (G) and time spent in center zone (H) of Pde11a-cre mice upon optogenetic stimulation of ChR2-EYFP or EGFP group, respectively.

**i,** Representative traces and statistical analysis of ChR2-EYFP or EGFP group in the elevated plus maze test of Pde11a-cre mice upon optogenetic stimulation of $ZIm^{Pde11a}$ neurons.

**j-l,** Representative traces (**j**) and statistical analysis (**k** and **l**) of ChR2-EYFP or EGFP group in the real-time place preference test of Pde11a-cre mice upon optogenetic stimulation. Preference index (**k**), $P$ = 0.7724. Preference to light (**l**), $P$ = 0.7072.

**m,** Schematics of novel object recognition test with a familiar object and a novel object.

**n,** The statistical analysis show the duration of investigation of Pde11a-cre mice in 5min upon optogenetic stimulation of ChR2-EYFP or EGFP group. EGFP group, **$P$ = 0.0013. ChR2-EYFP group, $P$ = 0.7777

One-way ANOVA for (**b-d**). Two-way ANOVA with Sidak's multiple comparisons test for (**g**), (**h**) and (**i**). Two-tailed paired t test for (**k**) and (**n**). Two-tailed unpaired t test for (**l**). Data are presented as mean ± SEM. Source data are provided as a Source Data file.

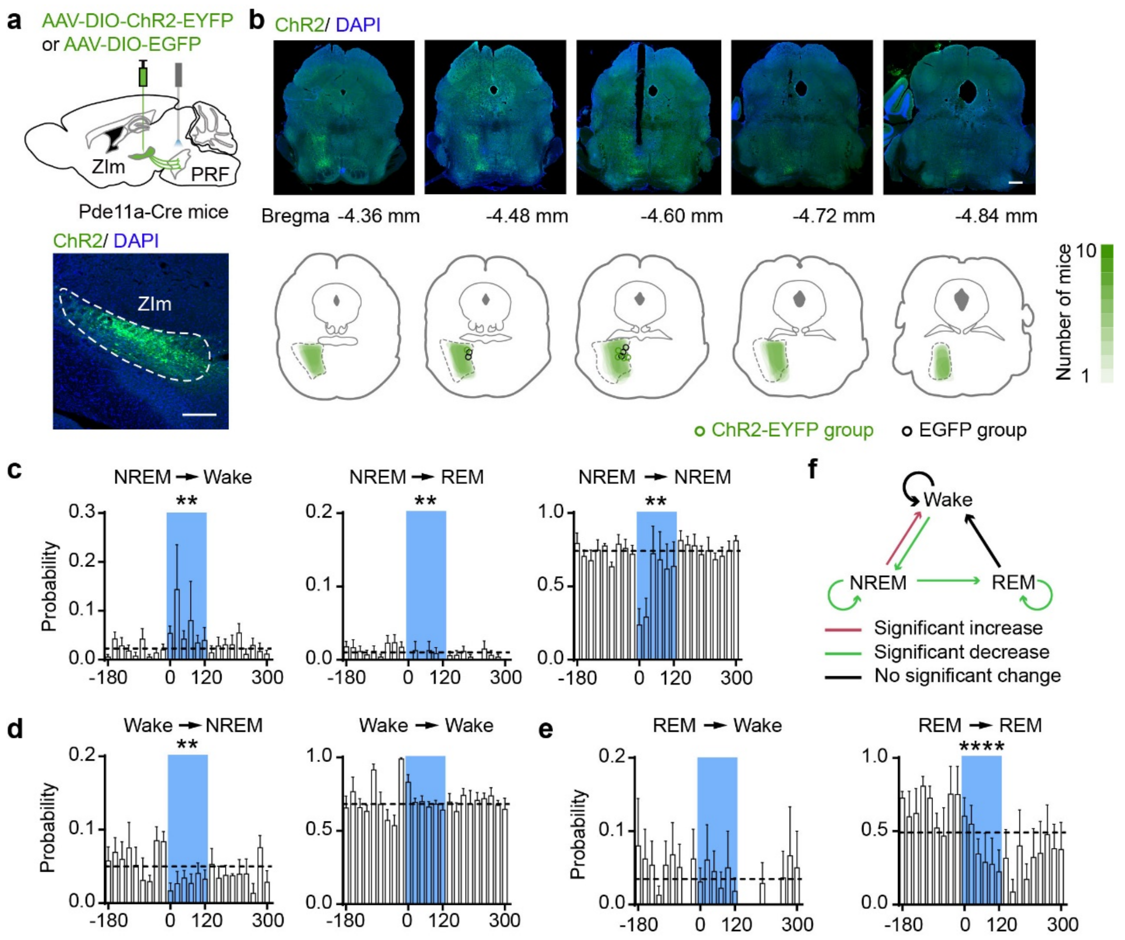


**Supplementary Figure 11: Optogenetic activation of ZIm$^{Pde11a}$ to PRF pathway significantly increases transition probabilities from sleep to wakefulness.**

**a,** Schematic diagram and representative images showing expression of ChR2-EYFP in ZIm of the Pde11a-cre mice. Scale bar, 250 μm. The experiment was independently repeated 5 times with similar results for each mouse strain.

**b,** The expression of ChR2-EYFP axon terminals of ZIm$^{Pde11a}$ neurons from anterior to posterior brain slice sections in PRF and schematic showing overlay of location of optical fiber placement in PRF with ChR2-EYFP or EGFP expression in ten mice after optogenetic stimulation was performed in PRF (lower). Green circles (ChR2-EYFP group, n = 5) and black circles (EGFP group, n = 5) indicate location of fiber placement for optogenetic stimulation in PRF. Scale bar, 500 μm. Green shaded areas indicate the expression range of neuronal terminals.

**c-e,** The transition probability between each pair of brain states upon optogenetic stimulation of the ZIm$^{Pde11a}$ to PRF pathway. Each bin indicates the transition probabilities within each 20 s period. Dashed line, baseline transition probability. Blue column indicates laser stimulation period (20 Hz, 120 s). $^{**}P$ < 0.05, $^{****}P$ < 0.0001, one-way ANOVA. n = 5 mice. Data are presented as mean ± SEM.

**f,** Schematic diagram showing transition probabilities between different brain states. Red, significant increase. Green, significant decrease. Black, no significant change. Source data are provided as a Source Data file.

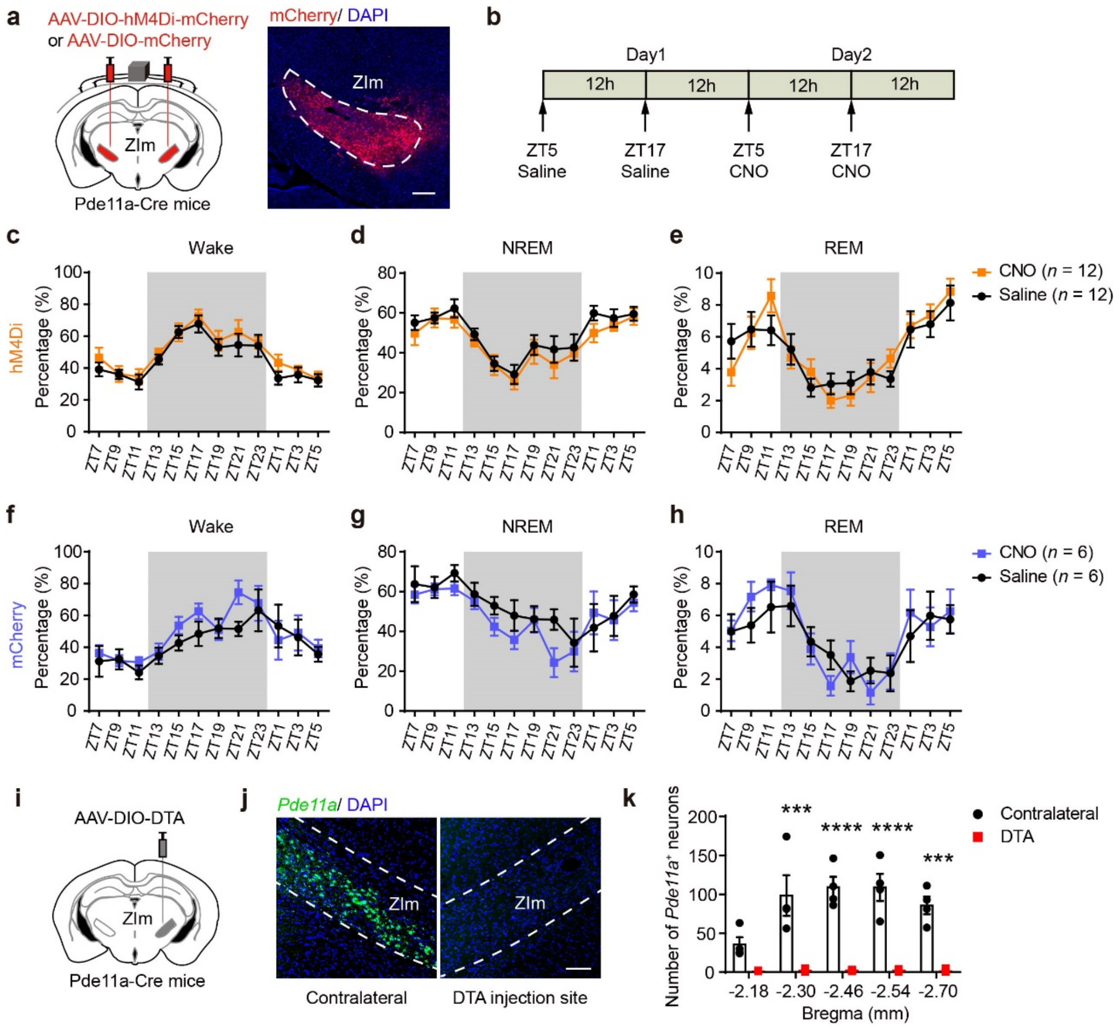


**Supplementary Figure 12: Chemogenetic inhibition of ZIm$^{Pde11a}$ neurons has little effects on sleep-wakefulness cycle.**

**a,** Schematic diagram and representative images showing the stereotaxic bilateral injection of AAV mediated Cre-dependent expression hM4Di-mcherry or mCherry (as control) into the medial ZI of Pde11a-Cre mice. Scale bar, 250 μm. The experiment was independently repeated 12 times with similar results for each mouse strain.

**b,** Schematic showing sleep recording conditions with saline- or CNO-injection in hM4Di-group and mCherry-group mice.

**c-h,** Percentage of time spent in wake, NREM sleep and REM sleep in 2-h bins following injection of saline or CNO at ZT5 and ZT17 during 24 hours in hM4Di-group (**c-e**) or mCherry-group mice (**f-h**). Grey bar indicates the dark phase (ZT12-ZT0).

**i-k,** Representative images (**j**) and statistical results (**k**) showing $Pde11a^{+}$ neuronal loss in the ZIm from mice with AAV-DIO-DTA unilateral injection in the ZIm. ***$P$ = 0.0002, ****$P$ < 0.0001, ***$P$ = 0.0007, n=4 mice. Scale bar, 100 μm. The experiment was independently repeated 4 times with similar results for each mouse strain.

Two-way ANOVA and Sidak's multiple comparisons test for (**c-h**) and (**k**). Data are presented as mean ± SEM. Source data are provided as a Source Data file.

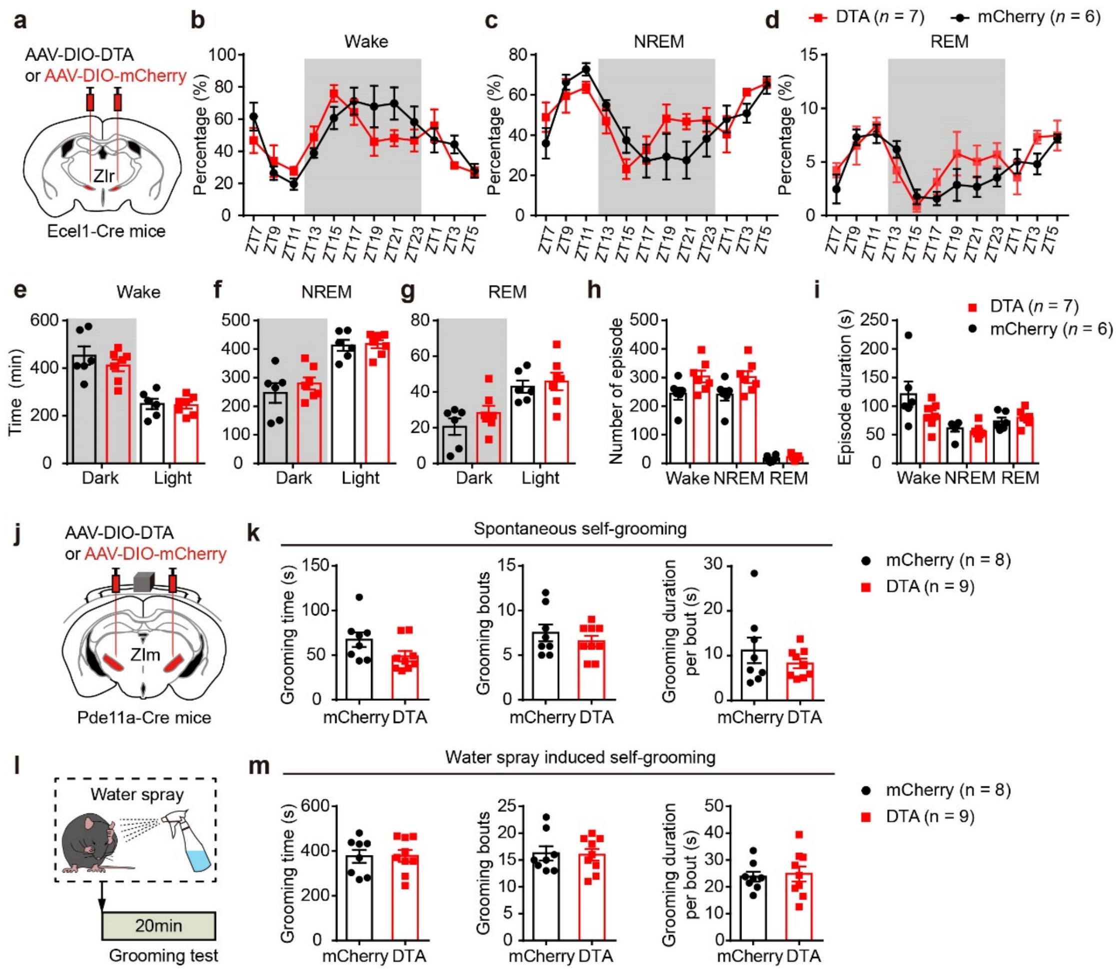


**Supplementary Figure 13: The ablation of ZIr$^{Ecel1}$ neurons has little effect on the sleep-wake cycle, and the ablation of ZIm$^{Pde11a}$ show no discernible effect on the self-grooming behavior.**

**a,** Schematic diagram showing the stereotaxic bilateral injection of AAV mediated Cre-dependent expression of DTA to ablate the ZIr$^{Ecel1}$ neurons of Ecel1-Cre mice.

**b-d,** Statistical analysis showing the percentage of time spent in wake, NREM sleep and REM sleep in 2-h bins in DTA-group mice compared with mCherry-group mice. Grey bar indicates the dark phase (ZT12-ZT0).

**e-g,** Time spent in wake (**e**), NREM sleep (**f**) and REM sleep (**g**) in dark or light phase across the 24-hour sleep-wake cycle in DTA-group mice compared with mCherry-group mice. Grey bar indicates the dark phase.

**h,** Number (**h**) and Duration (**i**) of wake, NREM sleep and REM sleep episodes during the dark phase.

**j,** Schematic diagram showing the stereotaxic bilateral injection of AAV mediated Cre-dependent expression of DTA to ablate the $ZIm^{Pde11a}$ neurons of Pde11a-Cre mice.

**k,** Statistic analysis of time spent for spontaneous self-grooming, grooming bouts and grooming duration per bout in 20 min in DTA-group mice compared with mCherry-group mice.

**l,** Schematic diagram showing the behavioral paradigm for measurement of water-spray induced self-grooming behavior.

**m,** Statistic analysis of water-spray induced self-grooming time, grooming bouts and grooming duration per bout in 20 min following water spray in DTA-group mice.

Two-way ANOVA and Sidak's multiple comparisons test for (**b-d**). Two-tailed unpaired t test for (**e-i**), (**k**) and (**m**). Data are presented as mean ± SEM. Source data are provided as a Source Data file.

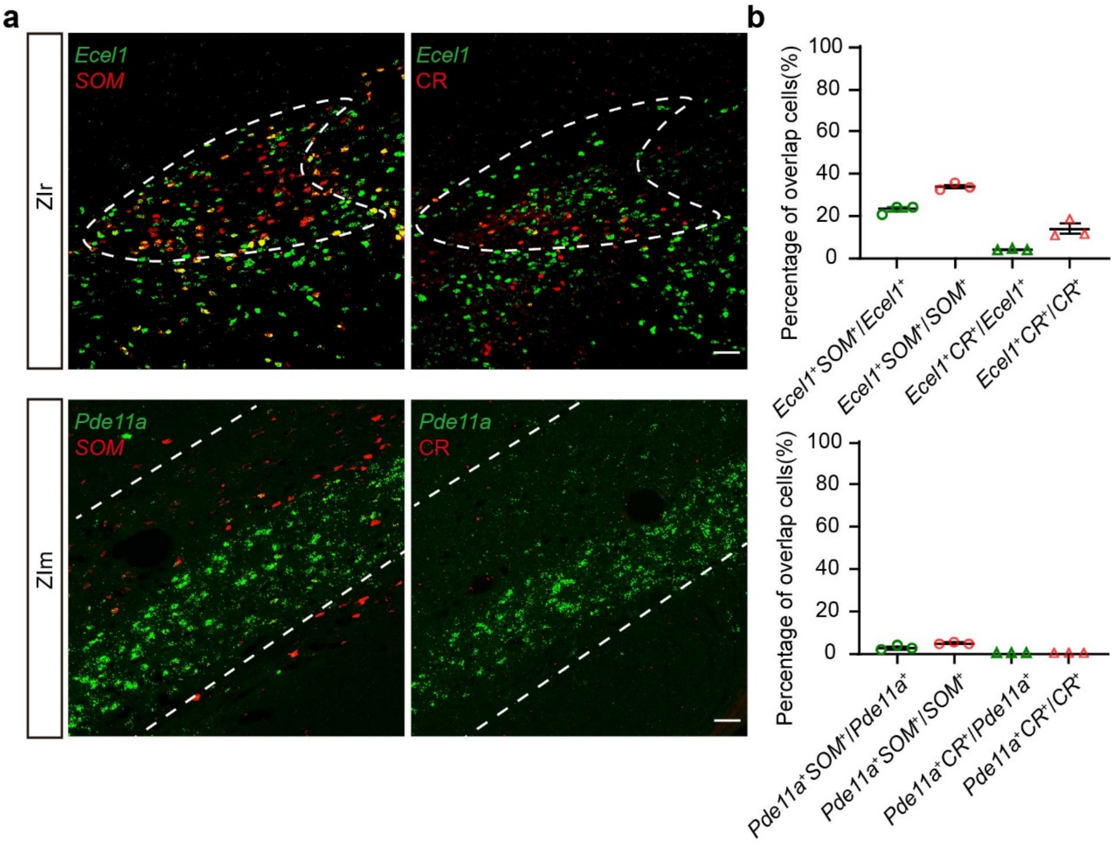


**Supplementary Figure 14: The co-expression level of *Ecel1* or *Pde11a* with *SST* and CR in ZI.**

**a,** Representative images of the FISH labeling of the mRNAs for *Ecel1*, *Pde11a* (green), *SST* (red) and proteins for CR (red) in the ZIr (upper) and ZIm (lower). Scale bars, 50μm. Upper: bregma, -1.20 mm. Lower: bregma, -2.40 mm. The experiment was independently repeated 3 times with similar results for each mouse strain.

**b,** Quantification of the proportion of *SST* and CR positive cells in ZIr *Ecel1* expressing (upper) and ZIm *Pde11a* expressing (lower) cells. Upper: the percentage of *Ecel1*$^+$*SST*$^+$ cells and *Ecel1*$^+$CR$^+$ cells in ZIr relative to all *Ecel1*$^+$, *SST*$^+$ or CR$^+$ cells in ZIr (*Ecel1*$^+$*SST*$^+$/*Ecel1*$^+$, 23.12 ± 1.12%; *Ecel1*$^+$*SST*$^+$/*SST*$^+$, 33.93 ± 0.96%; *Ecel1*$^+$CR$^+$/*Ecel1*$^+$, 4.10 ± 0.15%; *Ecel1*$^+$CR$^+$/CR$^+$, 13.80 ± 2.38%). Lower: the percentage of *Pde11a*$^+$*SST*$^+$ cells and *Pde11a*$^+$*CR*$^+$ cells in ZIm relative to all *Pde11a*$^+$, *SST*$^+$ or CR$^+$ cells in ZIm (*Pde11a*$^+$*SST*$^+$/ *Pde11a*$^+$, 2.58 ± 0.61%; *Pde11a*$^+$*SST*$^+$/*SST*$^+$, 4.83 ± 0.27%; *Pde11a*$^+$CR$^+$/*Pde11a*$^+$, 0.00 ± 0.00%; *Pde11a*$^+$CR$^+$/CR$^+$, 0.00 ± 0.00%). n = 3 mice (6 brain slices per mouse). Data are presented as mean ± SEM.

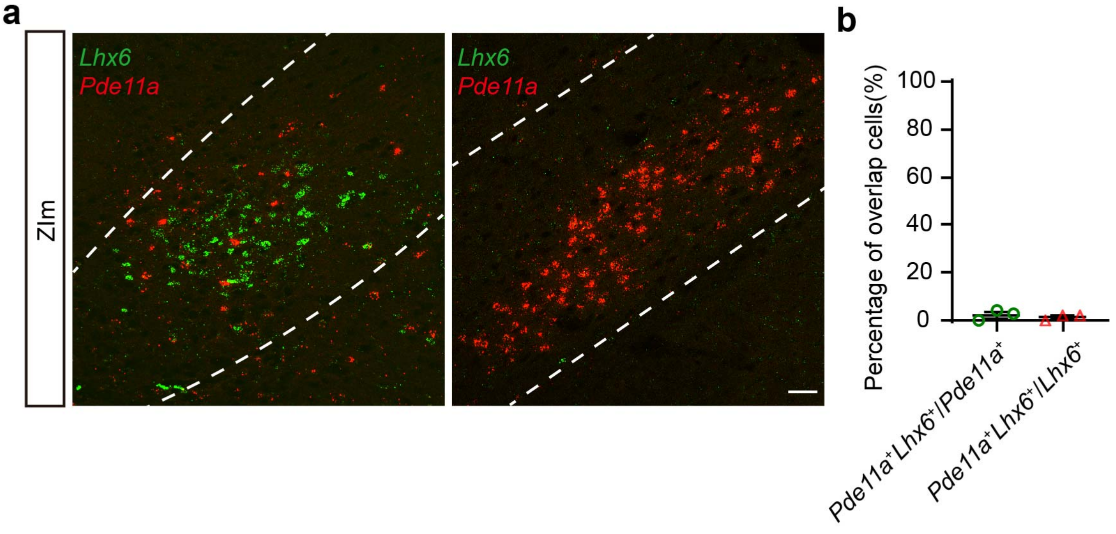


**Supplementary Figure 15: The expression and distribution of *Lhx6* and *Pde11a* in ZI.**

**a,** Representative images of the FISH labeling of the mRNAs for *Pde11a* (green), and *Lhx6* (red) in ZIm. Scale bars, 50μm. Left: bregma: -1.80mm; Right: bregma: -2.40mm. The experiment was independently repeated 3 times with similar results for each mouse strain.

**b,** Quantification of the percentage of *Pde11a* and *Lhx6* positive cells in ZIm. The percentage of *Pde11a*$^+$*Lhx6*$^+$ cells relative to all *Pde11a*$^+$, or *Lhx6*$^+$ cells in ZIm (*Pde11a*$^+$*Lhx6*$^+$/*Pde11a*$^+$, 2.23 ± 1.18%; *Pde11a*$^+$*Lhx6*$^+$/*Lhx6*$^+$, 1.38 ± 0.69%). N = 3 mice (6 brain slices). Data are presented as mean ± SEM.

**Supplementary table**

Table S1. Cluster marker gene list of ZI GABAergic neurons.

The genes differentially expressed in cells of subclusters were analyzed using FindMarkers function. The FindMarkers function used the two-sided Wilcoxon Rank-Sum test as its statistical method.
